# Characterizing the short-latency evoked response to intracortical microstimulation across a multi-electrode array

**DOI:** 10.1101/2021.11.22.469555

**Authors:** Joseph Sombeck, Juliet Heye, Karthik Kumaravelu, Stefan M. Goetz, Angel V. Peterchev, Warren M. Grill, Sliman Bensmaia, Lee E. Miller

## Abstract

**Objective:** Persons with tetraplegia can use brain-machine interfaces to make visually guided reaches with robotic arms. Without somatosensory feedback, these movements will likely be slow and imprecise, like those of persons who retain movement but have lost proprioception. Intracortical microstimulation (ICMS) has promise for providing artificial somatosensory feedback. If ICMS can mimic naturally occurring neural activity, afferent interfaces may be more informative and easier to learn than interfaces that evoke unnaturalistic activity. To develop such biomimetic stimulation patterns, it is important to characterize the responses of neurons to ICMS.

**Approach:** Using a Utah multi-electrode array, we recorded activity evoked by single pulses, and short (~0.2 s) and long (~4 s) trains of ICMS at a wide range of amplitudes and frequencies. As the electrical artifact caused by ICMS typically prevents recording for many milliseconds, we deployed a custom rapid-recovery amplifier with nonlinear gain to limit signal saturation on the stimulated electrode. Across all electrodes after stimulation, we removed the remaining slow return to baseline with acausal high-pass filtering of time-reversed recordings. With these techniques, we could record ~0.7 ms after stimulation offset even on the stimulated electrode.

**Main results:** We recorded likely transsynaptically-evoked activity as early as ~0.7 ms after single pulses of stimulation that was immediately followed by suppressed neural activity lasting 10–150 ms. Instead of this long-lasting inhibition, neurons increased their firing rates for ~100 ms after trains. During long trains, the evoked response on the stimulated electrode decayed rapidly while the response was maintained on non-stimulated channels.

**Significance:** The detailed description of the spatial and temporal response to ICMS can be used to better interpret results from experiments that probe circuit connectivity or function of cortical areas. These results can also contribute to the design of stimulation patterns to improve afferent interfaces for artificial sensory feedback.

## Introduction

Efferent brain-machine interfaces (BMIs) have advanced to the point where a spinal-cord injured patient can move a robotic arm using signals recorded from motor cortex (Collinger et al. 2013; Wodlinger et al. 2014; Hochberg et al. 2012). Without somatosensory feedback, the effectiveness of the movements generated through these interfaces will be limited, perhaps like those of people who have lost somatosensation (Ghez et al. 1990; Sainburg et al. 1995). Intracortical microstimulation (ICMS), which has been shown to elicit percepts in rats, monkeys, and humans (Devecioğlu and Güçlü 2017; Fridman et al. 2010; London et al. 2008; Romo et al. 2000), is a promising approach for providing artificial somatosensory feedback via an afferent interface (Tabot et al. 2013; Flesher et al. 2016). In the first such bidirectional BMI, monkeys could move a virtual arm to explore the “texture” of different virtual objects, a property conveyed by two different temporal patterns of ICMS (O’Doherty et al. 2011). The monkeys moved the arm sequentially to the objects to find the one with the rewarded texture. More advanced methods have been used to supply a spinal-cord injured patient with information about object contact location and force (Flesher et al. 2016; Flesher et al. 2021). Using a robotic arm, the patient was able to pick up, move, and place objects faster using vision combined with ICMS feedback than with visual feedback alone, primarily because they spent less time attempting to grasp the object (Flesher et al. 2021).

Restoring proprioception, the sense of position and movement of the body, has proven more difficult. In one approach, monkeys learned to reach to invisible targets using ICMS feedback through eight arbitrarily chosen electrodes which provided information about the error vector between hand and target position (Dadarlat, O’Doherty, and Sabes 2015). Monkeys only learned to use this feedback after a few months of training. To shorten this long learning period, it may be possible for ICMS to provide more naturalistic feedback (Bensmaia and Miller 2014). Our lab attempted to evoke perceptions of hand movement by stimulating on sets of four electrodes in somatosensory cortical area 2, that all had similar preferred directions (Tomlinson and Miller 2016). This biomimetic approach was successful for six of seven sets of electrodes in one monkey but failed in three other monkeys. The reason for the difference between monkeys remains unexplained, but had we been able to monitor evoked activity in each case, it may have been clearer.

To better interpret experiments which use ICMS and to achieve more successful mimicry of naturally occurring activity, it will likely be important to quantify the evoked response of neurons to a range of stimulus parameters. However, recording at short latency after stimulation is difficult due to the large shock artifact it causes (Hao, Riehle, and Brochier 2016; Weiss et al. 2018). Many experiments have been limited to recordings made on electrodes hundreds of microns away or even on a separate array (Hao, Riehle, and Brochier 2016; Butovas and Schwarz 2003; Chen et al. 2020; Allison-Walker et al. 2021), thereby missing evoked activity near the stimulated electrode. Further, previous studies have typically characterized the evoked response to only single pulses of stimulation, whereas future afferent interfaces will need to employ trains of stimulation throughout a grasp and/or movement (Flesher et al. 2021).

We developed novel hardware and software techniques allowing us to record ~0.7 ms after stimulation offset on every electrode in an implanted microelectrode array, including even the stimulated electrode. We characterized the neural responses to single pulses, short trains (~0.2 s), and long trains (~4 s) of stimulation at different amplitudes and frequencies. Consistent with other studies, we observed excitatory activity evoked in neurons immediately after stimulation with a single pulse followed by long periods of inhibition (Hao, Riehle, and Brochier 2016; Butovas et al. 2006; Butovas and Schwarz 2003). In contrast, after short, high-frequency trains of stimulation, neurons greatly increased their firing rates for ~0.1 s. During long trains, the excitatory response recorded on the stimulated electrode decayed, while the response on non-stimulated electrodes was typically maintained throughout the train. The results in this paper can inform the interpretation and design of stimulation patterns for providing somatosensory feedback.

## Methods

### Animal Subjects

We performed experiments using two male rhesus macaques. Monkey H was 12.0 kg and monkey D was 10.0 kg when we performed the experiments. We performed all procedures in this study in accordance with the Guide for the Care and Use of Laboratory Animals. The institutional animal care and use committee of Northwestern University approved all procedures in this study under protocol #IS00000367.

### Implant and data collection

Each monkey was implanted with a 96-electrode sputtered iridium-oxide multi-electrode array with 1.0 mm electrodes (Blackrock Microsystems, Salt Lake City, UT) in the proximal arm area of somatosensory cortical area 2. In addition to surface landmarks, we recorded intraoperatively from the cortical surface while manipulating the arm and hand to find the arm representation (for more details, see Weber et al. 2011). We performed sensory mappings after implantation to confirm that recorded neurons had receptive fields corresponding to the proximal arm.

We used the Blackrock Stim Headstage, Front-End amplifier, and Neural Signal Processor (Blackrock Neurotech, Salt Lake City, UT) to record signals at 30 kHz. We delivered ICMS from the Blackrock CereStim R96. Unless otherwise noted, electrodes were stimulated with biphasic pulses, each phase lasting 200 μs and separated by 53 μs. We used the sync line from the CereStim R96 to determine stimulation onset.

During all experiments, monkeys performed a center-out reaching task while holding the handle of a robotic manipulandum (for more details, see (London and Miller 2012)) or sat idly in the chair. Stimulation was delivered independently of the monkey’s behavior.

### Pipeline to record at short latencies after ICMS

Typically, ICMS causes large electrical artifacts which prevent neural recordings for an extended period after stimulation. When using the Blackrock Stim Headstage and Front-End amplifier to record on the stimulated electrode, the recorded signal saturated the amplifier for several milliseconds (Fig. 1a, dashed lines), after which the signal slowly recovered to baseline. To enable recording at shorter latencies, we developed a rapid-recovery amplifier (RRA, see Supplementary Materials) and used it instead of the Blackrock Stim Headstage and Front- end Amplifier. The RRA has several features that allow it to operate on the same electrode as the stimulator, yet still recover rapidly after stimulation. The wide input range (± 15 V) of the first stage of the RRA prevents input voltage clamping and current shunting as well as output saturation during the stimulus pulse. To prevent saturation of subsequent stages, the gain of the RRA declines rapidly from a maximum of ~1000 to a minimum of 1 during large dynamic swings of the front-end voltage. The output of the RRA, which was limited to ± 5 V, was connected to an analog input on the Blackrock Neural Signal Processor (Fig. 1b).

**Fig. 1.**
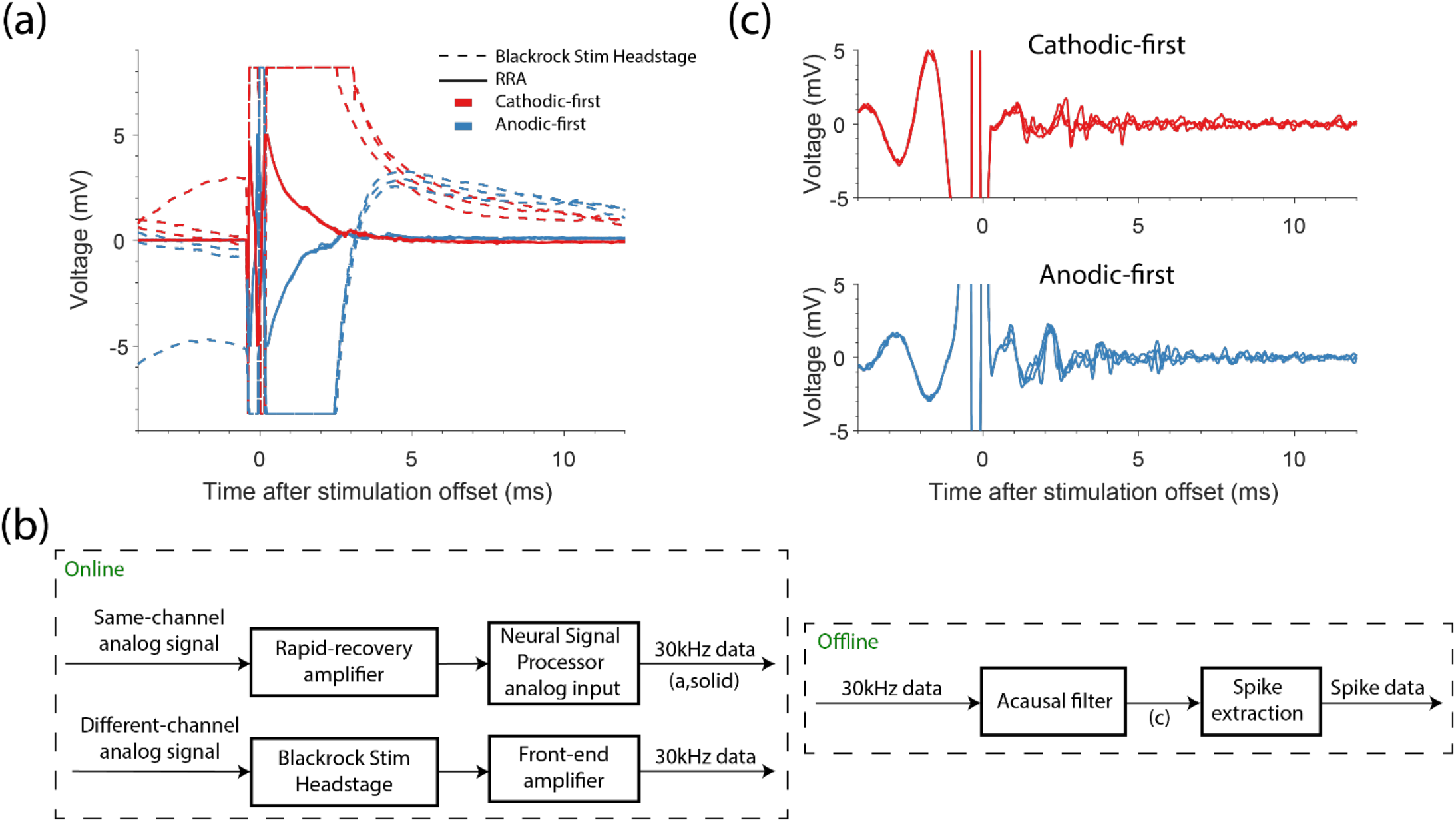
Overview of artifact reduction pipeline. (a) Example recordings from the stimulated channel are shown when recording with the Blackrock Stim Headstage and Front-end amplifier (dashed lines) and the rapid recovery amplifier (RRA; solid lines). We stimulated with anodic-first (blue) or cathodic-first (red) biphasic pulses with phase duration of 200 μs, phases separated by 53 μs, and with an amplitude of 50 µA. (b) Block diagram depicting the artifact reduction pipeline. The rapid-recovery amplifier receives signals and passes them to the Blackrock Neural Signal Processor. Signals from channels that were not stimulated were amplified by the Blackrock Stim Headstage and Front-End amplifier. All signals were sampled at 30 kHz and filtered offline. After filtering, we extracted spikes and then sorted the spike data. (c) Voltages recorded using the RRA after acausal time-reversed high-pass filtering. Traces in (c) correspond to those in (a).

To measure the progressive gain recovery of the RRA after stimulation when stimulating and recording on the same electrode, we monitored the size of the artifact evoked by much lower current stimulation on a remote electrode. We tested gain recovery following alternating cathodic- and anodic-first biphasic pulses at 10 Hz, with amplitudes of 5–30 μA in 5 μA steps and 40–100 μA in 10 μA steps. We tested 25 stimulation electrodes across the two monkeys and delivered 32 ± 2 (mean ± sd) pulses per condition. The remote channel was stimulated at 3000 Hz for 4.5 ms, with cathodic-first biphasic pulses (53 μs pulse length with 53 μs between phases). We used 1 μA to monitor gain recovery on four stimulation electrodes in one session, and 5 μA on the remote channel in later sessions.

Even with the RRA, full recovery to baseline took ~3 ms (Fig. 1a, solid lines). While a high-pass filter removed this drift, ringing caused by filtering the large artifact prevented neural recording for ~10 ms. Instead, we applied a 500 Hz high-pass Butterworth filter acausally, backwards in time, thereby preventing the introduction of a ringing artifact (Fig. 1c). We adjusted the timestamps of recorded spikes to account for the ~100 μs phase shift caused by filtering. Even with this acausal filtering we avoided filtering through the artifact, which would have obscured the pre-stimulus data (as seen in Fig. 1c). To account for the changing gain of the RRA, we divided the recorded signal by the measured gain recovery. After filtering, we extracted neural activity by finding threshold crossings and then sorting single units using OfflineSorter (Plexon Inc., Dallas, TX).

Recordings on non-stimulated electrodes using the Stim Headstage and Front-end Amplifier were saturated for ~0.7 ms after stimulation offset. In our testing, the RRA did not shorten the recording latency on non-stimulated electrodes. Because of this, we did not use the RRA when recording on non-stimulated channels. Nevertheless, we filtered acausally before extracting neural activity as we did for recordings made on the stimulated electrode.

### Evaluating the performance of the rapid-recovery amplifier and acausal, time-reversed filtering

To evaluate the performance of our pipeline for recording neural activity on the stimulated electrode, we tested how well we could recover simulated activity at different latencies after stimulation. We recorded electrical artifacts on 10 different stimulated electrodes across two monkeys with the RRA or with the Stim Headstage and Front-End amplifier. To simulate neural activity, we recorded naturally occurring spike waveforms during a period without stimulation and then summed these waveforms with the recorded artifacts at random times after stimulation. We added spike waveforms at random latencies between 0.2–7 ms following 50% of the stimuli for each of the 10 electrodes. For each electrode, we generated 200,000 stimulation artifacts, half from recordings made with the RRA and half with the standard Blackrock hardware. We tested the same amplitudes described above for measuring RRA gain but used only cathodic-first pulses since our subsequent experiments used this polarity. We computed the percentage of spikes recovered by comparing the time stamps of recovered spikes to the artificial ones, tolerating ± 0.33 ms of error.

### Stimulation protocol for characterizing the evoked response

After evaluating our recording capability, we characterized the response evoked on the stimulated channel by single pulses or pulse trains. Table 1 shows the numbers of sessions for each monkey, electrodes tested, neurons recorded, and trains per condition for all experiments. The final column (inter-train period) indicates the time between successive stimulation conditions. We slightly jittered the inter-train period for each condition to prevent synchronizing stimulation with any physiological process by adding 0–100 ms sampled from a uniform distribution.

**Table 1.** Experimental parameters are shown for the single pulse and short train experiments, continuous (Cont.) long train experiments when recording on either the stimulated channel (stim) or non-stimulated channels (nonstim), and intermittent (Inter.) long train experiments when stimulating at 179 Hz or 131 Hz. The number of sessions for monkey H and monkey D are denoted with ‘H’ and ‘D’ respectively.

| | # sessions | # stimulation electrodes | # neurons | # pulses or trains (mean ± std.) | Inter-train period (s) |
| --- | --- | --- | --- | --- | --- |
| **Single Pulse** | 4H; 4D | 29 | 30 | 82.6 ± 14.4 | 0.5 |
| **Short Train** | 7H; 5D | 19 | 19 | 253 ± 17.2 | 0.5 |
| **Cont. Long Train (stim)** | 6H; 4D | 24 | 25 | 8 ± 0 | 20 |
| **Cont. Long Train (nonstim)** | 6H; 4D | 24 | 437 | 8 ± 0 | 20 |
| **Inter. Long Train (179 Hz)** | 4H; 4D | 22 | 22 | 8 ± 0 | 20 |
| **Inter. Long Train (131 Hz)** | 2H; 2D | 12 | 12 | 8 ± 0 | 20 |

In initial experiments we measured the response evoked by single pulses at a range of amplitudes. In four experiments, we tested 10–60 μA in 10 μA steps, 80 and 100 μA. Later, we probed the lower stimulation amplitudes more thoroughly using 10–30 μA in 5 μA steps and 40, 50, and 100 μA for another 4 electrodes, then added a 5 μA condition for the final 21 electrodes.

We next characterized the response to short (~0.2 s) trains. We stimulated at 50 μA and at 20, 49, or 94 Hz for 7 neurons. After noticing a modest decay in the response of neurons throughout the 0.2 s train at 94 Hz, we added a 179 Hz condition for the remaining 12 neurons. After short, high frequency trains, we observed rebound excitation, which we analyzed with this data.

We then characterized the evoked response to longer (~4 s) trains of stimulation, a duration that approximates that required for a BCI user to grasp and move an object (Flesher et al. 2021). Because the recorded neural response decayed rapidly with 179 Hz stimulation, we used a maximum of 131 Hz when stimulating with 4-s long trains. We stimulated with all combinations of 20, 40, 60 μA and 51, 80, 104, 131 Hz. Data were collected simultaneously on the stimulated and non-stimulated channels during this experiment. The results for non-stimulated channels may include a given neuron activated by different stimulation electrodes.

### Data analysis

All data analysis was performed using MATLAB (MathWorks Inc., Natick, MA). To quantify the amount of activity evoked by each pulse, we counted spikes between 0.5 and 5.0 ms after the offset of each pulse and averaged across pulses. To account for different baseline firing rates across neurons, we subtracted the expected number of spontaneous spikes based on the baseline firing rate measured 10 ms to 80 ms before onset of single pulses or 0.2 to 2 s before train onset.

We computed an activation threshold for each neuron in response to single pulses. To do so, we measured the proportion of stimulation pulses with at least one spike occurring 0.5–5 ms after stimulation offset for each condition and neuron. We defined the activation threshold as the smallest amplitude at which the proportion of trials with a spike was significantly larger than that expected based on the baseline firing rate (Chi-Square test, α < 0.05). We determined if a neuron was responsive to long trains of stimulation in a similar manner. Since the evoked response decayed throughout long trains, we considered only the first 20 pulses in each 8 trains. For each condition, neurons with significantly more spikes than chance (Chi-Square test, α < 0.05) were considered responsive.

Multiple spikes were typically evoked at consistent latencies by single stimulus pulses. We grouped spikes based on their response latency across trials for each neuron and condition. To do so, we convolved the spike train after stimulation offset with a non-causal Gaussian kernel with width of 0.2 ms, then averaged across pulses. We found peaks in this average with MATLAB’s *findpeaks* algorithm. This algorithm uses “prominence”, the height of a peak and its location relative to other peaks, to measure how much a peak stands out (for more details, see Supplementary Materials for more details). We required peaks to have a minimum prominence of 1.0 and to be separated by at least 0.7 ms. This algorithm also computes the width at half maximum of each peak. Spikes that occurred within the width of each peak were included in the corresponding group. We measured the latency of each peak and computed the standard deviation of the spike times within each group. Our results were only slightly affected by small changes to the smoothing kernel width, minimum peak spacing, and minimum prominence.

After an evoked response, many neurons underwent either long-lasting inhibition or rebound excitation, which we quantified by computing the average firing rate across trials using a two-bin running average across 5 ms bins. We defined an inhibitory response as firing rates below three-quarters of the mean baseline firing rate for two consecutive bins (a similar threshold as (Butovas and Schwarz 2003)) and measured the time the firing rate remained below this threshold. We defined a rebound excitatory response if two consecutive bins exceeded twice the mean baseline firing rate and the corresponding duration.

For many neurons, the evoked response decreased throughout long trains of stimulation. We measured the decay rate for each responsive neuron. To do so, we measured the mean firing rate in 50 ms bins from 0.0 to 3.9 s after train onset, accounting for the stimulation artifact by removing 1 ms of time per stimulation pulse in each bin. We then fit the firing rate with an exponential decaying function,

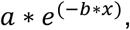

with *a* as the intercept and *b* as the decay rate. During intermittent stimulation, we only included bins which contained at least 2 stimulation pulses.

### Statistical Analysis

Statistical analyses were performed using MATLAB (MathWorks Inc., Natick, MA). We used linear and logistic models to analyze many of our results. We included two interaction terms in the model when analyzing the effect of amplitude, time, and amplifier on the proportion of simulated spikes recovered: one between amplitude and amplifier, to test whether the effect of amplitude was reduced with the RRA, and a second between time and amplifier, to see if the rate of spike recovery increased with the RRA. When analyzing the effect of amplitude on the latency of evoked spikes, we included an interaction term between amplitude and spike group number. Finally, we included an interaction term between amplitude and frequency when analyzing the decay rate throughout long trains of stimulation. After fitting the models, we performed F-tests on the resulting parameters from the linear models and t-tests on the resulting parameters from the logistic models.

We performed Wilcoxon rank-sum tests to compare the magnitude of evoked activity recorded on non-stimulated channels at 20 μA and 60 μA for each neuron and stimulation electrode pair. Here, we aggregated data across stimulation frequencies. We used this same test for differences between the decay rates for intermittent and continuous stimulation for each duty cycle. We aggregated data across durations, which did not have a large effect.

## Results

### Recording pipeline performance

To evaluate the performance of the RRA, we first measured its dynamic gain recovery after stimulation at different amplitudes. We delivered a single biphasic pulse through the electrode to which the RRA was connected and simultaneously injected a known signal to a remote electrode (Fig. 2a). After acausal, time-reversed filtering, we determined the gain of the amplifier by dividing the amplitude of each pulse in the known signal by the mean amplitude of the final three pulses, which were well after full gain recovery. The mean gain recovery curves, aggregated for 25 stimulation electrodes across two monkeys, are shown for both cathodic- and anodic-first pulses at several stimulation amplitudes in Fig. 2b. We compared the gain of the amplifier at 1 ms across stimulation amplitudes and polarities using a repeated measures ANOVA (F(26,481) = 40.6, p = 6.58E-104). The gain of the amplifier recovered more slowly as amplitude increased (F(1,481) = 762.65, p = 2.8×10^−101^) and roughly 140 μs faster for cathodic-first pulses than for anodic-first pulses (F(1,481) = 142.2, p = 6.7×10^−29^). Subsequently, when measuring actual neural signals, we accounted for the changing gain by dividing the recorded signal by the gain function (Fig. 2a, bottom).

**Fig. 2.**
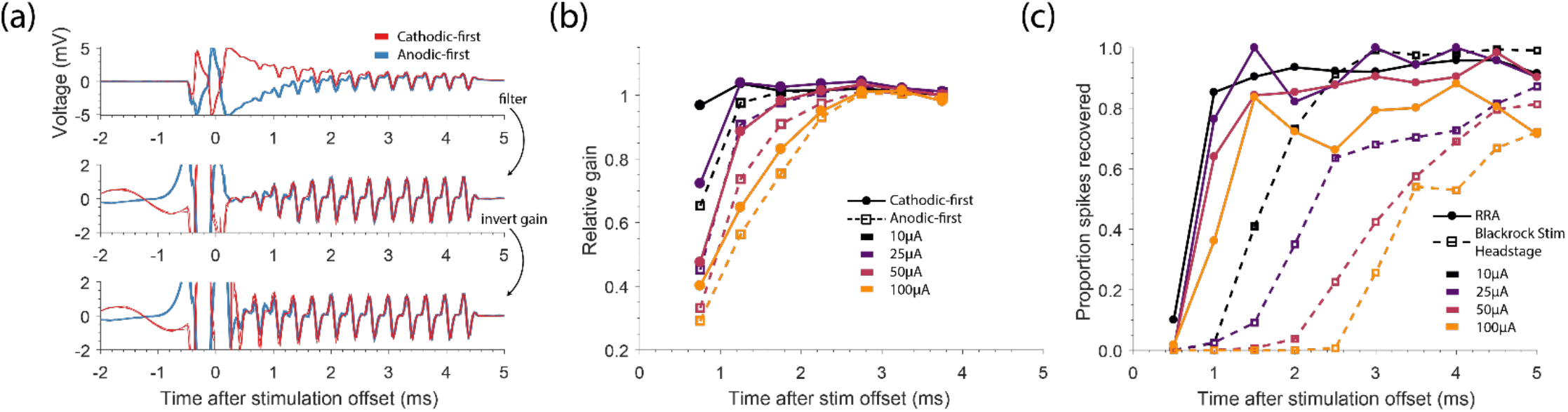
Evaluation of rapid-recovery amplifier (RRA). (a) Example recordings on the stimulated channel when evaluating the gain of the RRA are shown both before (top) and after (middle) acausal, time-reversed filtering, and after accounting for the changing gain (bottom). We stimulated with biphasic anodic-first (blue) or cathodic-first (red) pulses with phase duration of 200 μs, phases separated by 53 μs, and an amplitude of 50 μA. Pulses were simultaneously delivered on a remote channel to inject a ‘known’ signal. (b) The relative gain of the RRA for stimulation at different amplitudes. The gain was determined by measuring the peak-to-peak voltage of the injected signal. (c) Spikes were artificially added to artifact traces recorded on the stimulated channel. The proportion of simulated spikes recovered using our pipeline for both the RRA (solid lines) and the Blackrock Stim Headstage (dashed lines) across stimulation amplitudes.

We tested the ability to recover spikes following stimulation by adding representative, naturally occurring spike waveforms to actual recordings of stimulation artifacts to establish a ground-truth reference. The proportion of these spikes that could be recovered with the Blackrock Headstage and with the RRA are shown in Fig. 2c. We used logistic regression to predict the proportion of spikes recovered based on the stimulation amplitude and time after stimulation, (overall model χ^2^ = 1.97×10^3^, p ≅ 0). Not surprisingly, spike recovery worsened with increasing stimulation amplitude regardless of amplifier (p = 5.8×10^−46^, t-test), but spikes were recovered at much shorter latencies with the RRA than with the Blackrock Stim Headstage (p = 3.7×10^−31^, t-test). The RRA also reduced the effect of amplitude (p = 0.0015, t-test) and increased the recovery rate (p = 0.0090, t-test).

### Excitatory and inhibitory response to single pulses of ICMS

After evaluating the performance of the RRA, we used it for a series of experiments to quantify the neural responses evoked on the stimulated electrode. We first characterized the excitatory and inhibitory responses following single pulses across a wide range of current amplitudes (5–100 μA). Responses for an example neuron are shown in Fig. 3a. While it was not possible to record throughout stimulation (red shading indicates region obscured by the artifact), using the RRA allowed us to record many spikes that we could not have seen if we had used the Blackrock Headstage (grey shading). The number of spikes evoked above baseline firing across amplitudes is shown in Fig. 3b. The number increased significantly as amplitude increased (overall model F(30,223) = 4.88, p =1.36×10^−12^; amplitude factor F(1,223) = 12.029, p = 6.3×10^−4^). Among the 29 out of 30 neurons that were activated with the range of currents tested, the median activation threshold was 10 μA (Fig. 3c).

**Fig. 3.**
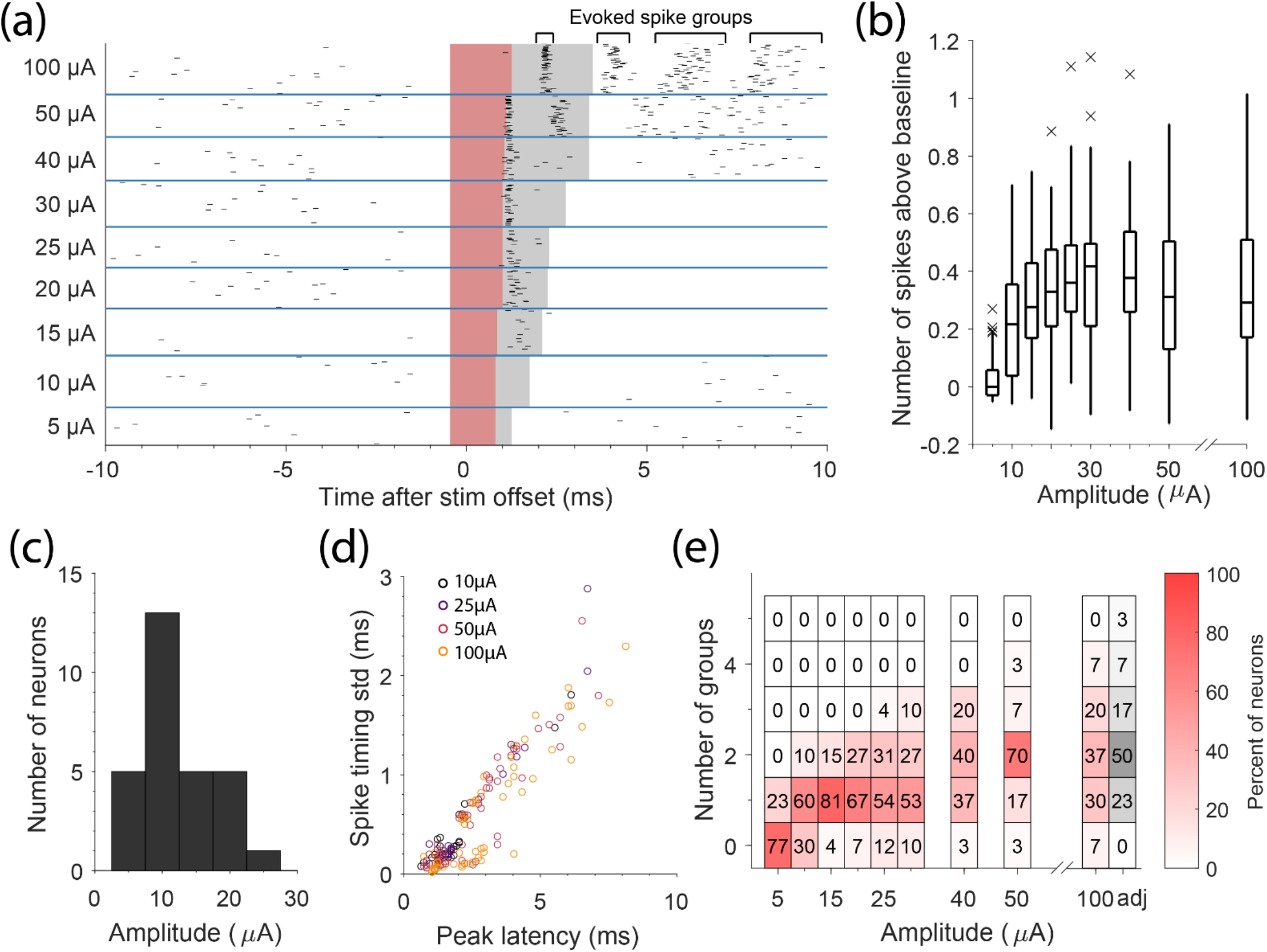
Excitatory response on the stimulated channel after single pulses of stimulation. (a) Response of an example neuron recorded on the stimulated channel in response to single cathodic-first pulses at different amplitudes. Each row is a different stimulation trial, and each tick represents an action potential from this neuron. Blue, horizontal lines separate stimulation trials at different amplitudes. Red shading depicts the time interval while we were unable to record neural signal with the RRA. Grey shading depicts the corresponding time if we had used the Blackrock Stim Headstage and Front-End amplifier. (b) The number of evoked spikes above baseline is shown across neurons for each stimulation amplitude. X’s mark outliers. (c) Distribution of activation thresholds across neurons. (d) The standard deviation of spike times within an evoked spike group is shown against the latency of that group for different stimulation amplitudes. (e) The number of groups evoked for different stimulation amplitudes. The number within each box and the shading of each box indicates the percentage of neurons. When stimulating with 100 μA, the extended artifact likely obscured the entire initial group of spikes. When we determined that this occurred, we increased the number of groups by one. The adjusted numbers are displayed in grey (‘adj’).

Sufficiently high stimulation amplitude evoked multiple spikes within 10 ms of stimulation offset. These spikes occurred at consistent latencies across trials, with later spikes having more varied timing than earlier ones. To quantify this, we grouped evoked spikes based on their latency (Fig. 3a and Supplementary Materials show example groups). Fig. 3d shows the standard deviation of spike times within a group compared to the latency of that group for multiple stimulation amplitudes. This standard deviation increased significantly as group latency increased (overall model F(32,302) = 103, p =1.0×10^−142^; latency factor F(1,302) = 574.13, p = 8.1×10^−72^). We also noticed that latencies decreased as stimulation amplitude increased, seen as a leftward shift in Fig 3a as current increased to 100 μA, at which point the artifact likely obscured the first recorded group of evoked spikes.

Using a linear model across all neurons, we determined that the latency of groups decreased by 3.6 ± 0.7 μs/μA as amplitude increased (overall model F(31,303) = 55.1, p =7.3×10^−106^; amplitude factor F(1,303) = 25.497, p = 7.7×10^−7^).

The number of spike groups evoked across stimulation amplitudes is shown in Fig 3e with the color and text within each box representing the percentage of neurons. At 100 μA, the extended artifact and decreased latency likely obscured the entire initial group of spikes. When we determined that this occurred, we increased the number of groups by one. Even without this compensation, the number of groups increased significantly with stimulation amplitude (overall model F(20,223) = 5.65, p = 4.9×10^−15^; amplitude factor F(1,223) = 85.5, p = 1.9×10^−17^).

After stimulation, neuronal activity was typically suppressed for ~10–150 ms depending on the stimulation amplitude (Fig. 4a). We fit a linear model to predict inhibition duration by amplitude across neurons (overall model F(30,180) = 1.9, p=0.0057) and found that increasing stimulation amplitude significantly increased the inhibition duration (amplitude factor F(1,180) = 32.43, p = 5.0×10^−8^) and increased the fraction of cells undergoing inhibition (Fig. 4b). Stimulation amplitudes ≥ 40 μA caused inhibition in ~90% of neurons.

**Fig. 4.**
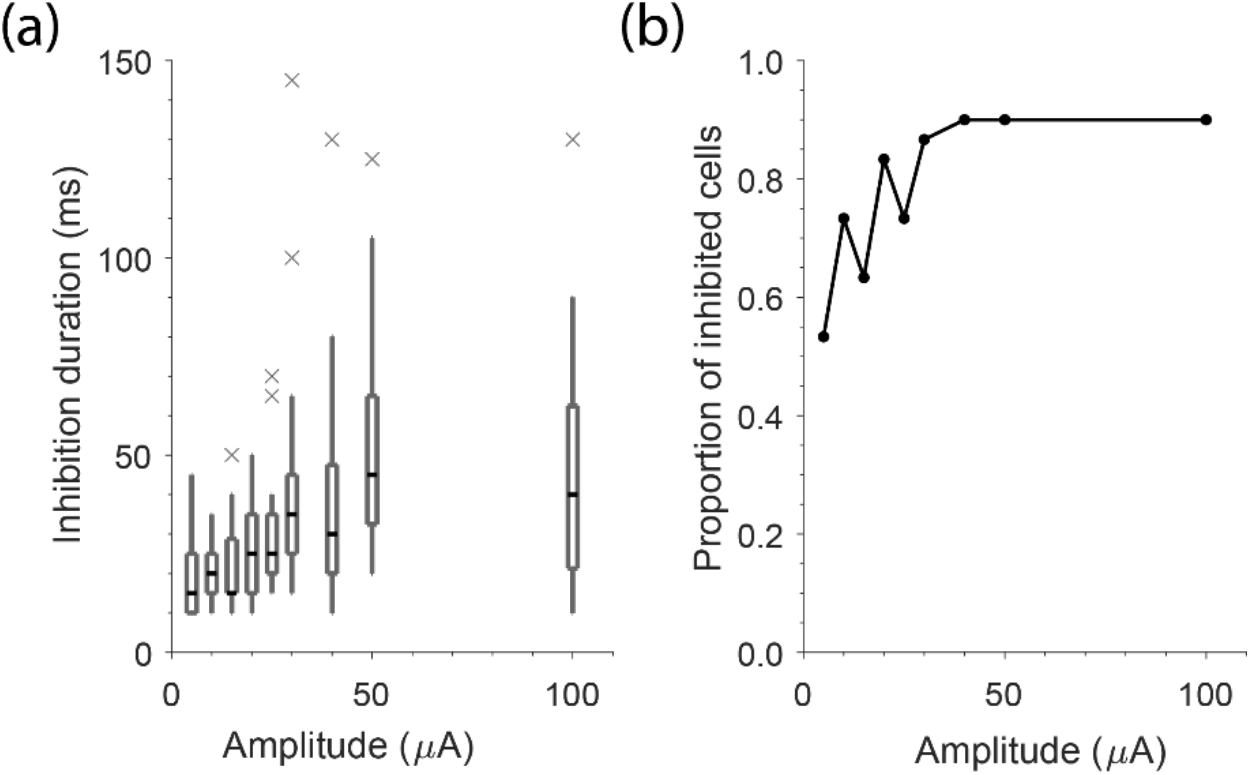
Inhibitory response recorded on the stimulated channel after single pulses of stimulation. (a) The inhibition duration across neurons recorded on the stimulated channel after single cathodic-first pulses of stimulation across stimulation amplitudes. (b) The fraction of cells with an inhibitory response is shown for each stimulation amplitude.

### Temporal response to trains of ICMS

We hypothesized that the activity evoked by ICMS would decrease throughout long stimulus trains as a consequence of the long-lasting inhibition on stimulated electrodes following single pulses (Fig 4). To test this, we stimulated on single electrodes with 4-s long trains at several amplitudes (20, 40, 60 μA) and frequencies (51, 80, 104, 131 Hz). The mean responses across eight trains for nine of the 12 stimulation conditions are shown as grey traces in Fig. 5a for an example neuron. For this neuron, the evoked response rapidly decayed throughout the train, particularly for the larger amplitudes and frequencies. For the 21.5 ± 2.0 neurons that were activated significantly for each condition (Chi-Square test, α < 0.05), we computed a decay rate by fitting the firing rate during stimulation with an exponential (Fig. 5b). Using a linear model (F(26,231) = 14.7, p = 8.6×10^−36^), we determined that the evoked response decayed significantly faster with greater stimulation amplitude or frequency (amplitude: F(1,231) = 119, p = 9.4×10^−23^; frequency: F(1,231) = 134, p = 8.8×10^−25^). Increased frequency (amplitude) had a larger effect at higher amplitudes (frequencies) (interaction term: F(1,231) = 71.4, p = 3.3×10^−15^).

**Fig. 5.**
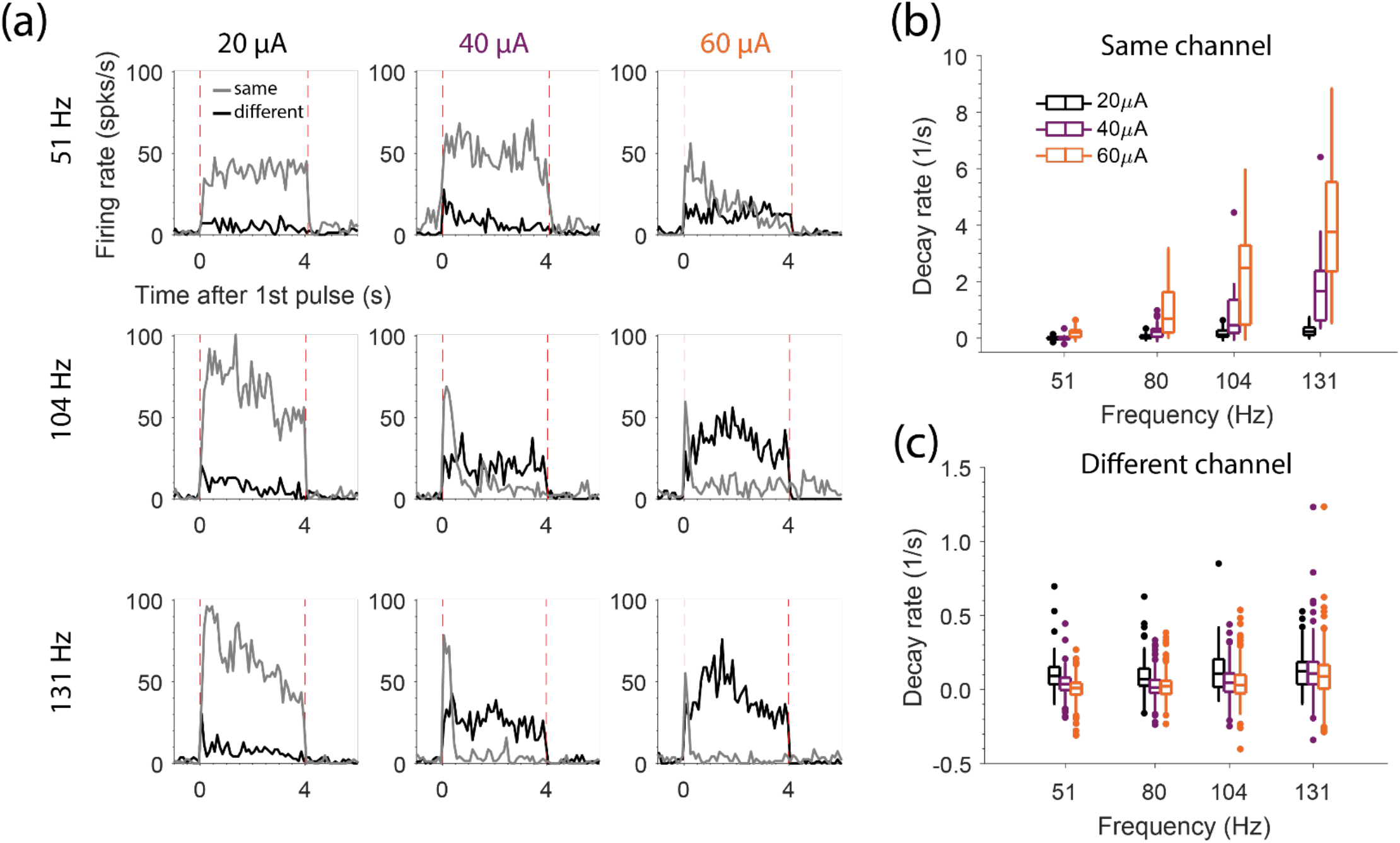
Decay rate throughout 4-s long trains of stimulation. (a) The mean firing rate across stimulation trials for the same neuron when the channel it was recorded on was stimulated (grey) and when a different channel was stimulated (black) for different stimulation amplitudes (columns) and frequencies (rows). Amplitudes and frequencies are noted above and to the right of the panels, respectively. Vertical, red dashed lines indicate train onset and offset. (b) The decay rates across neurons recorded on the stimulated channel for each amplitude and frequency. Points indicate outliers. (c) The decay rates for each neuron recorded on non-stimulated channels for each amplitude and frequency. Note the smaller y-limits in (c) compared to (b).

Since the response on the stimulated channel decays rapidly, we wondered whether allowing recovery time during the stimulus train would reduce the decay rate. To test this, we turned stimulation on and off throughout the train (see Supplementary Materials for more information). However, intermittent stimulation did not significantly change the decay rate compared to continuous stimulation at frequencies chosen to match the number of pulses (Wilcoxon rank-sum test, p > 0.05 for all). This shows that the effect of intermittent stimulation is similar to simply reducing the mean stimulus frequency.

If neurons recorded on non-stimulated electrodes were driven transsynaptically by neurons activated near the stimulated electrode, then we would expect to see a similar rapid decay in the evoked activity for neurons on non-stimulated electrodes. If, on the other hand, neurons even on distant electrodes are driven directly, their decay rate may differ from that of neurons recorded on the stimulated electrode. To determine this, we examined the neuronal activity evoked on non-stimulated electrodes. In contrast to the response on the stimulated electrode, this activity did not decay appreciably (Fig. 5a, black traces). The maintained response of all 437 neuron/stimulation electrode combinations was shown here was essentially like this example (Fig. 5c). Using a linear model with data aggregated across amplitudes and frequencies (F(81,3269) = 51.6, p ≅ 0), we determined that the evoked response decayed significantly faster for neurons recorded on the stimulated channel than on non-stimulated channels (F(1,3269) = 980.3, p = 2.0×10^−188^). These results imply that the response on non-stimulated electrodes is driven directly, or by evoked activity that occurs before we can record it.

After the end of an ICMS train, we expected neurons on the stimulated electrode to be inhibited for many milliseconds, as we observed with single pulses (Fig. 4). Indeed, low-frequency, 50 μA trains delivered for ~0.2 s caused inhibition (see example in Fig. 6a) in about 50% of neurons, lasting from 10-250 ms (Fig. 6b). Faster stimulus frequency increased inhibition duration (Model: F(19,21) = 6.38, p = 5.9×10^−5^, frequency factor F(1,21) = 36.0, p=6.0×10^−6^) but this effect was not observed in all 16 tested neurons. At 179 Hz, the highest frequency we tested, the fraction of cells with an inhibitory response was only ~8%. Instead of inhibition in these cases, we observed a large burst of activity immediately after the stimulation train. This rebound excitation occurred for 75% of cells following stimulation at 179 Hz and lasted from ~25-240 ms (Fig. 6c). If a neuron exhibited rebound excitation for multiple stimulation frequencies, higher frequencies almost always resulted in longer lasting rebound. During the longer 4-s trains, we observed rebound excitation very infrequently (2/25 cells) potentially because of the longer train duration.

**Fig. 6.**
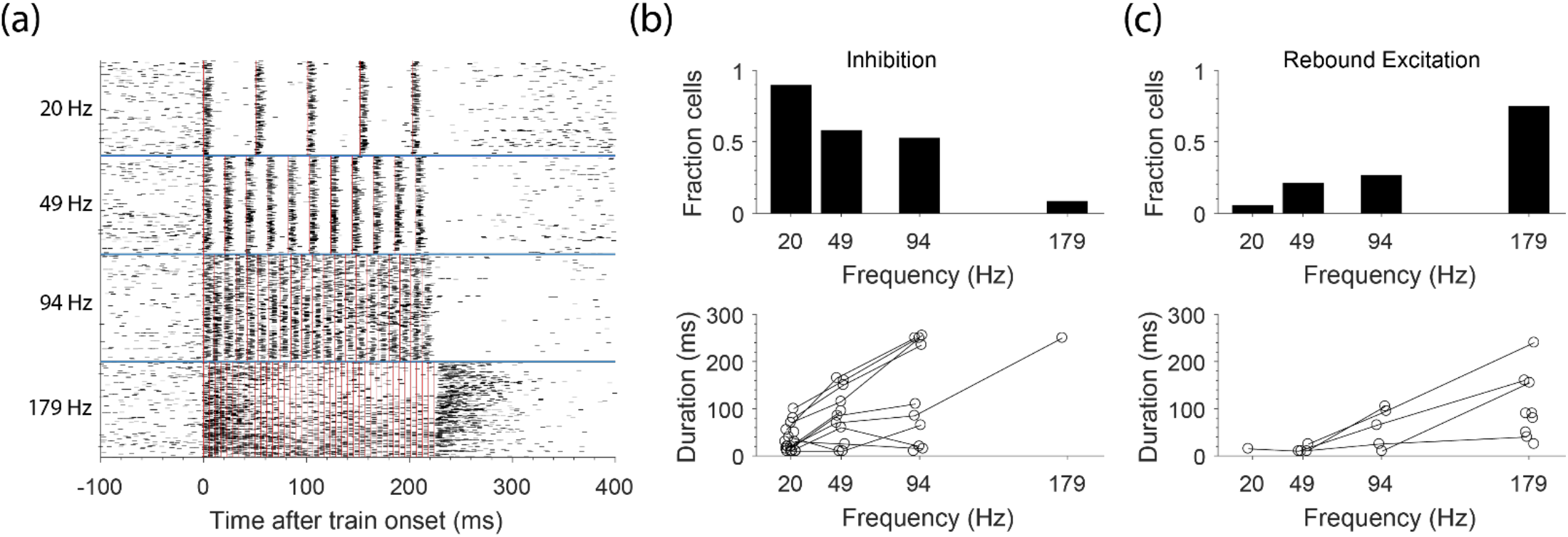
Rebound excitation recorded on the stimulated electrode after short trains of stimulation. (a) The response of an example neuron recorded on the stimulated electrode during ~200 ms trains at different frequencies. Red lines indicate stimulation pulses. Stimulation frequencies are shown on the left of the figure for 50 μA stimulation. (b) The fraction of cells that displayed an inhibitory response after the end of the short trains (top) and the duration of the inhibitory responses (bottom) for each frequency. (c) The fraction of cells that displayed rebound excitation (top) and the duration of the rebound excitation (bottom) for each frequency. Lines connecting points represent data from the same neuron.

### Spatial pattern of the response to ICMS trains

Both increased amplitude and frequency typically increase ICMS detectability, perhaps because of increased charge delivery (Kim et al. 2015; Otto, Rousche, and Kipke 2005; Sombeck and Miller 2019). Increasing amplitude leads both to more activity near the stimulated electrode (Fig. 3) as well as a wider spread of activity recorded across a multi-electrode array (Hao, Riehle, and Brochier 2016; Stoney, Thompson, and Asanuma 1968; Kumaravelu et al. 2021), likely because increased amplitude results in more charge delivered per pulse. Greater frequency, though, does not change the charge per pulse and thus may not lead to equivalent effects. To study these effects, we measured activity on non-stimulated electrodes throughout 4-s trains of continuous stimulation. Fig 7a shows the increase in firing rate above baseline of each neuron for each stimulation electrode aggregated across two monkeys against distance from the stimulated electrode. We computed the mean firing rate above baseline for each neuron and amplitude / frequency combination repeated across 8 trains (Fig. 7b). For each stimulation electrode, we only analyzed neurons that had activation thresholds at or below 20 μA when stimulating at 51 Hz. We normalized the responses by the number of pulses in the train and pooled them across stimulation frequencies. The evoked activity per pulse at 60 μA was significantly larger than that at 20 μA for 290 out of 437 neurons (p<0.001, Wilcoxon rank-sum test). Using a linear model (F(125,1362) = 23.5, p = 8.9×10^−260^), we determined that increasing amplitude increased the evoked firing rate per pulse (F(1,1362) = 1165.5, p = 4.4×10^−185^). Increasing frequency also significantly increased the evoked activity per pulse (F(1,1362) = 11.1, p = 0.00089), an effect that was, however, two orders of magnitude smaller than that of increasing amplitude. In addition to this small effect of each pulse, increased frequency increased overall evoked activity to a greater extent because of the greater number of pulses in four seconds.

**Fig. 7.**
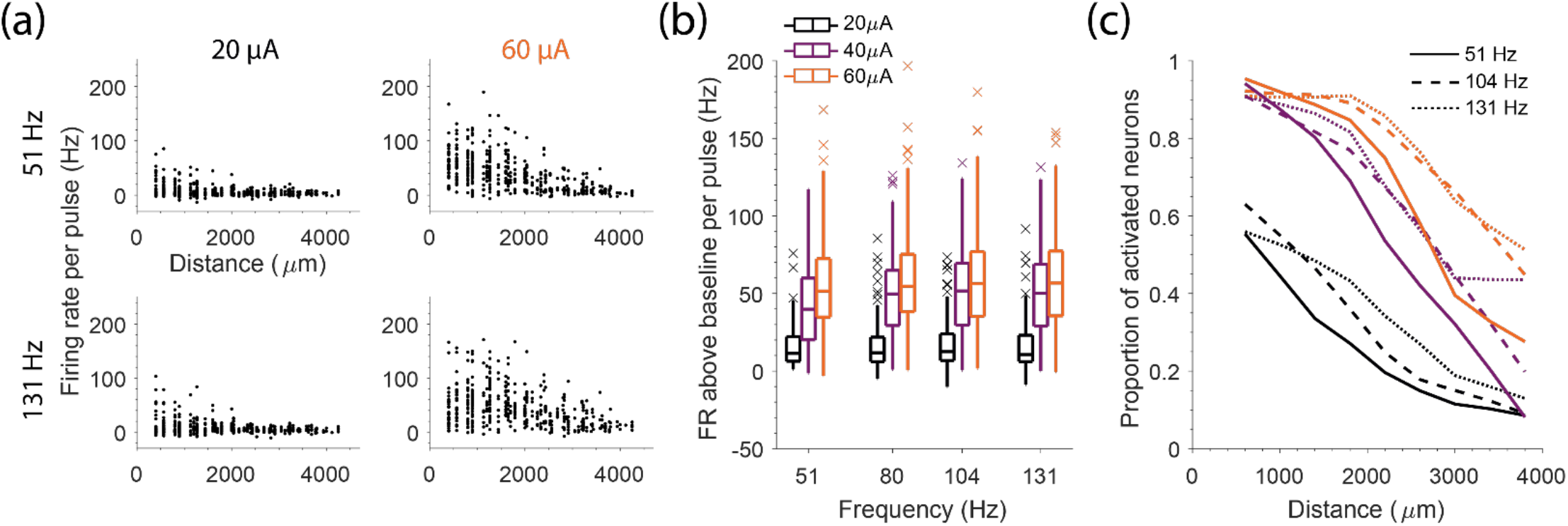
Evoked response on non-stimulated channels during 4-s long trains of ICMS. (a) The firing rate above baseline against distance from the stimulated electrode for different amplitudes (columns) and frequencies (rows). Each point represents a neuron and stimulated electrode pair. (b) The firing rate above baseline per pulse for each frequency and amplitude condition for responsive neurons. X’s mark outliers. (c) The proportion of neurons activated at different distances is shown for a subset of amplitudes (color) and frequencies (line-style).

We also hypothesized that increased stimulation amplitude would increase the distance at which neurons are activated while increased frequency would not. Data for a subset of stimulation conditions are shown in Fig. 7c. We used logistic regression to determine the effect of amplitude, frequency, and distance on the proportion of activated neurons (overall model χ^2^(104) = 1.69×10^3^, p ≅ 0). While increasing either amplitude or frequency increased the proportion of activated neurons (amplitude: p = 8.9×10^−159^; frequency: p = 1.6×10^−9^), the effect of frequency was an order of magnitude smaller.

## Discussion

We developed hardware and software tools to enable recording at short latency after ICMS. With these tools, we were able to record roughly 0.7 ms after the end of stimulation, even from the stimulated channel. We investigated the evoked response to single pulses, short trains, and long trains of ICMS of varying amplitude and frequency to better understand the neural response to stimulation. Here, we compare our methods and results to those of previous studies, discuss the mode of activation for the spikes we recorded, and how our results may impact the sensations evoked by ICMS in afferent interfaces.

### Comparison of artifact suppression to previous techniques

Recording neurophysiological potentials immediately after passing current through an electrode is difficult; the large shock artifact typically prevents recordings for many milliseconds. We developed and evaluated a rapid-recovery amplifier (RRA) to enable short latency recordings, particularly on the stimulated electrode. The RRA clamps the voltage below that which would saturate downstream electronics by reducing gain as the magnitude of the input voltage increases (Fig. 2b). An alternative approach to shorten the duration of the artifact is to electrically disconnect the recording system during stimulation (Zhou, Johnson, and Muller 2018). While this approach is effective on non-stimulated electrodes (Hao, Riehle, and Brochier 2016), it cannot remove artifact on the stimulated electrode, which is caused by residual polarization of the electrode itself (Venkatraman et al. 2008). Our approach is similar to clamping the slew rate (first derivative) of a signal, as has been done previously (Epstein 1995). By reducing the gain, we reduced the size of the artifact and prevented saturation, thereby allowing us to record at earlier latencies. Another important advantage of the RRA is the wide input voltage range (±15 V) that avoids input clamping and stimulus current shunting of the relatively high voltage (< 10 V) stimulus pulses. A benefit of our approach is that the RRA can be placed in front of pre-existing recording systems, in our case, the Cerebus system from Blackrock Neurotech. Saturation can also be prevented by using an amplifier with a lower gain and/or an amplifier with a higher maximum input voltage (Jung, Kim, and Nam 2018; Rolston, Gross, and Potter 2009).

While the RRA prevents amplifier saturation that would otherwise be caused by the large shock artifact, the recorded signal still returns slowly to baseline after stimulation (Fig. 1a). This slow return is likely caused by slow dissipation of the residual charge on the electrode (Zhou, Johnson, and Muller 2018). To remove excess charge more quickly, custom electronics could be designed to actively discharge the electrode to a pre-stimulus voltage (Brown et al. 2008; DeMichele and Troyk 2003; Freeman 1971), although this may introduce switching artifacts that diminish the effectiveness of this approach.

The slow return to baseline can also be removed offline. When done with a high-pass filter, it is important not to filter through the shock artifact, as this can cause ringing and obscure the neural signal. One solution is to filter data beginning a fixed time after the end of stimulation (Hao, Riehle, and Brochier 2016). Instead, we filtered acausally, backwards in time so that any ringing would be pushed before the stimulation, leaving the post-stimulus data clean (such acausally displaced ringing can be seen before 0 ms in Fig. 1c). This approach does not require defining a time at which the shock artifact has ended, though it does push neural signal back in time ~100 μs. We compensated for this time shift by adjusting the time stamps of recorded spikes by 100 μs. With the RRA and acausal, time-reversed filtering, we were able to record ~0.7 ms after stimulation offset, even on the stimulated electrode (Fig. 2c), revealing spikes that we could not have recorded with the Blackrock Stim Headstage (grey shading in Fig. 3a).

### Mode of activation of recorded spikes on stimulated and non-stimulated channels

ICMS can evoke action potentials both directly and transsynaptically (Tehovnik et al. 2006). Directly evoked spikes occur because stimulation changes the membrane potential of cells near the electrode, causing them to fire. Action potentials are typically initiated in axons, which have a higher density of sodium channels than do somas, resulting in lower activation thresholds (Nowak and Bullier 1998a, 1998b; Tehovnik et al. 2006). Action potentials then propagate antidromically to the cell bodies and orthodromically to presynaptic terminals, where they may elicit further activity transsynaptically.

We wondered whether the spikes we recorded on the stimulated electrode were evoked directly, at either the axon or soma, or transsynaptically. Since we have no direct way of testing this, we inferred the mode of activation from the latency of evoked spikes. We expect directly evoked spikes to be generated within 0.3 ms of the end of the cathodic phase (Gustafsson and Jankowska 1976; Jankowska, Padel, and Tanaka 1975; Stoney, Thompson, and Asanuma 1968), though we may actually observe these spikes somewhat later since they need to propagate from the site of initiation back to the soma. We estimated this potential antidromic distance and latency by first estimating how far spike initiation could have occurred from the stimulated electrode. To do so, we used Stoney’s square-root relationship (Stoney, Thompson, and Asanuma 1968):

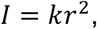

With *k* =1292 μA/mm^2^ and *I* = 10 μA, the median activation threshold of neurons in our study (Fig. 3b) the maximum spike initiation distance is ~100 μm. Since somas can be recorded up to ~150 μm from the recording electrode (Maynard, Nordhausen, and Normann 1997), the maximum distance an action potential could travel before being recorded is ~250 μm. With a propagation speed of 1 μm/μs (Swadlow 1990), the maximum latency at which we expect to see a directly evoked spike is 0.55 ms after the end of the cathodic phase, (0.3 ms after the end of our biphasic pulses). Hence, the earliest spikes we were able to see on the stimulated electrode (0.7 ms; Fig. 3c), were undoubtedly transsynaptic.

We asked the same questions about the spikes recorded on non-stimulated electrodes. Due to the increased distance that evoked spikes could propagate, the latency at which we could record directly evoked spikes would also increase. For electrodes within 400 μm of the stimulated electrode, the maximum distance an action potential could travel is ~650 μm, making the longest directly evoked spike latency ~0.7 ms. This is very close to the earliest spikes we saw. For these immediately adjacent electrodes, it remains likely we are recording transsynaptic activation. For electrodes farther than 400 μm from the stimulated electrode, the maximum latency of direct activation overlaps with our recording latency, suggesting that some spikes may have been directly evoked.

Knowing that we inevitably missed some early, directly evoked spikes, we estimated the proportion of spikes this represented. Since we can begin recording ~0.7 ms after stimulation offset, we might miss at most one spike per pulse. In the worst case, where the neuron is directly activated by each pulse, 1.2 – 1.4 spikes are evoked per neuron across amplitudes (Fig. 3b). The average of 0.2 – 0.4 transsynaptically evoked spikes we recorded account for 17 – 30% of evoked spikes. Even though distance and amplitude affect the proportion of pulses which directly evoke a spike (Stoney, Thompson, and Asanuma 1968), these likely still make up a large proportion of evoked spikes near the stimulated electrode.

### Optical recording to measure directly evoked activity

While we were able to record on the stimulated electrode at a short latency after stimulation, we likely missed the earliest directly evoked activity occurring during the stimulus pulse. To record during the stimulus pulse, calcium imaging or voltage-sensitive dye imaging can be used, as these imaging methods are not affected by the shock artifact (Histed, Bonin, and Reid 2009). With optical methods, researchers have measured the spatial and temporal response of neurons during stimulation at various amplitudes, frequencies, and with different pulse shapes (Histed, Bonin, and Reid 2009; Michelson et al. 2018; Stieger et al. 2020). While optical methods have many advantages, they do come with some limitations. Currently, they cannot be used in humans, limiting their usefulness for brain-machine interfaces. Optical recordings also have lower temporal resolution than electrical recordings, with frames recorded at about 30 Hz (Histed, Bonin, and Reid 2009; Michelson et al. 2018). While optical imaging can be used with awake, behaving animals (Dombeck et al. 2007; Greenberg, Houweling, and Kerr 2008; Andermann, Kerlin, and Reid 2010), studies recording the evoked response to stimulation have typically used anesthesia, which affects the response properties of neurons, including the balance of excitation and inhibition (Histed, Bonin, and Reid 2009; Stieger et al. 2020; Tehovnik and Slocum 2013; Michelson et al. 2018; Tanaka et al. 2019).

### Qualitatively similar evoked responses are observed across different experimental conditions

Across many studies using different levels of anesthesia, animal models, and recording techniques, the evoked response to ICMS is qualitatively similar. After stimulation, neurons exhibit short-latency excitation due to direct or transsynaptic activation (Margalit and Slovin 2018; Tehovnik et al. 2006). We observed an increase in the amount of evoked activity from activated neurons and an increase in the spread of evoked activity with increasing amplitude (Fig. 3), consistent with previous observations (Hao, Riehle, and Brochier 2016; Butovas and Schwarz 2003). With increased frequency, we observed a small increase in the amount of evoked activity per pulse, an effect that is further amplified by the increased number of pulses (Fig. 7). After short-latency excitation, neural activity is typically suppressed for long periods, an effect likely mediated by GABA_B_ receptors (Butovas et al. 2006). The duration of this long-lasting inhibition increased with amplitude in our study (Fig. 4), in contrast to previous observations of neurons recorded farther from the stimulated electrode (Butovas and Schwarz 2003). After inhibition, we often saw a large increase in firing rate (Fig. 6) (Butovas and Schwarz 2003). This rebound excitation may be due to recurrent excitation within cortical circuits, mediated by calcium channels (Molineux et al. 2006; McElvain et al. 2010). Throughout trains of stimulation, we observed a rapid decay of the evoked activity recorded near the stimulated electrode (Fig. 5), similar to previous observations (Michelson et al. 2018). This rapid decay could be caused by the local activation of inhibitory neurons (Overstreet, Klein, and Helms Tillery 2013).

### Linking evoked activity to sensation

In monkeys, stimulation in tactile cortices evokes sensations at locations corresponding to the receptive field of neurons recorded on the stimulated electrode (Tabot et al. 2013). Different temporal patterns of stimulation can be distinguished and used to convey useful information (Hughes et al. 2021; Callier et al. 2020; Berg et al. 2013; London et al. 2008; Romo et al. 1998; Dadarlat, O’Doherty, and Sabes 2015). Similar observations have been made in humans with tetraplegia and neuropathy (Salas et al. 2018; Chandrasekaran et al. 2021; Fifer et al. 2020), including the ability of one person to identify which of multiple fingers of a robotic hand, linked to somatosensory cortex (S1) stimulation, were touched (Flesher et al. 2016). More recently, somatosensory ICMS was used to provide contact and pressure-related feedback, which improved their ability to control a robotic arm to reach and grasp (Flesher et al. 2021). The stimulus parameters in this most recent example were quite simple, a linear mapping from index and middle finger joint torques to appropriate electrodes. To improve the feedback provided by ICMS, it is important to understand the effect stimulation parameters, including amplitude, frequency, pulse width, and train length have on the quality and intensity of the evoked percept. Characterizing the sensations evoked by interacting electrodes would be very time consuming for patients (if even possible) given the very large stimulation parameter space. Instead, it may be possible to infer aspects of the evoked sensation indirectly, by recording evoked activity, an approach that may allow for more rapid testing of stimulation parameters for many electrodes.

The intensity of an evoked sensation can be increased by increasing any of these parameters, within limits (Kim et al. 2015; Otto, Rousche, and Kipke 2005; Sombeck and Miller 2019). However, amplitude and frequency have different effects on the resulting sensation, as shown by the different effects these parameters had on monkeys’ perceptions of noisy moving dot fields. At small amplitudes, stimulation biased the monkeys’ perception of the dot motion, in a direction predicted by the preferred directions of recorded neurons and with an effect size dependent on stimulus current (Salzman et al. 1992). However, near 80 μA, the monkeys’ performance returned to chance level, perhaps because of the recruitment of more distant neurons with less homogenous properties (Murasugi, Salzman, and Newsome 1993). In contrast, increasing frequencies as high as 500 Hz increasingly biased the monkeys’ perception. In our data, increasing amplitude greatly increased the spread of activation (Fig. 7). While increasing frequency also increased the spread of activation, this effect was much smaller than the increase in firing rate of those activated neurons. Our results are quite consistent with these behavioral observations.

Modulating frequency affects the nature of the evoked sensation. Monkeys can discriminate different frequencies of ICMS due to changes in either the intensity or quality of sensation (Romo et al. 1998; Berg et al. 2013; London et al. 2008). To distinguish between these possibilities, monkeys were trained to discriminate ICMS frequency when the amplitude of the stimulus train varied independently (Callier et al. 2020). Their ability to discriminate ICMS frequency independently of intensity differed greatly from electrode to electrode, implying that modulating frequency changed the quality of sensation for some (but not all) electrodes. Similar results were obtained with human patients, where modulating frequency sometimes changed the reported sensation (Hughes et al. 2021). In our data, the evoked response for some neurons decayed rapidly throughout trains of stimulation, while the response for others was maintained (Fig. 5). As frequency increases, some neurons will stop responding to stimulation throughout the train, changing the activated population of neurons and potentially changing the nature of the evoked sensation. To relate the temporal response of neurons to the quality of sensation, researchers would need to record both the evoked response and the reported quality of sensation evoked by stimulation through the same electrodes.

Surprisingly, human patients reported a decrease in sensation intensity with increasing frequency for some electrodes (Hughes et al. 2021), in contrast to the experiments in monkeys where increasing frequency increased the detectability of stimulation (Kim et al. 2015) and shortened the reaction time to stimulation (Sombeck and Miller 2019). We and others recorded a rapid decay in the amount of evoked activity on the stimulated electrode with time (Fig. 5 and Fig. 6a) (Michelson et al. 2018). Assuming these neurons contribute to the overall sensation intensity, intensity would be largest near the beginning and decrease throughout the train. As frequency increases, the intensity at the onset of the train likely increases due to the increased number of pulses, resulting in a more detectable signal and shorter reaction times. At high frequencies, though, the amount of evoked activity decays more rapidly, possibly leading to fewer total evoked spikes and a lower overall sensation intensity compared to stimulation at lower frequencies.

After the end of a reach or when an object is no longer grasped, the feedback provided by ICMS should end. We observed a large burst of activity after the end of high frequency trains (Fig. 6). This rebound excitation could potentially lead to sensations that persist beyond the end of the train, presumably extending the sensation. Since rebound excitation primarily occurred at high stimulation frequencies, it may be that there is a maximum frequency that future afferent interfaces can use to avoid this effect.

### Online recording in the presence of stimulation artifact

For most applications, afferent interfaces would only be useful when combined with an efferent interface, thereby providing both restored somatosensation and movement (Collinger et al. 2013; Flesher et al. 2016; O’Doherty et al. 2011). However, stimulation in S1 produces large artifacts in recordings from motor cortex (M1). With causal filters, neural signals can be recorded from M1 in a human ~0.7 ms after offset of stimulation applied in S1. At low stimulation frequencies, losing the ability to record from M1 for short periods after each pulse will not have much of an impact on decoding performance. When intended cursor velocity was decoded from M1, artificially dropping a random 20% of M1 signals caused only a 10% decrease in decoder performance (Fig. 8 in (Young et al. 2018)). While acausal, time-reversed filtering may allow for slightly earlier recordings, the increased amount of data would likely have a negligible impact on decoding performance.

However, as stimulation protocols become more complicated, with stimulation at high rates and on many electrodes (Bensmaia and Miller 2014; Sombeck and Miller 2019), the percentage of time in which signals can be recorded from M1 will decrease, further decreasing decoder performance. Stimulation at 333 Hz, either on a single electrode or across electrodes, would result in 50% loss of signal, assuming a total blanking duration of 1.5 ms per pulse (Weiss et al. 2018). With some non-trivial amplifier modifications to increase somewhat, the gain during the stimulus artifact, the RRA could potentially enable neural recordings even during the stimulus pulse, albeit at a significantly reduced gain. Although we did not explore them here, there are numerous approaches that could be used to extract neural signal from the artifact if the recorded signal is not saturated: adaptive filtering (Mendrela et al. 2016; Nag et al. 2015), template subtraction (Montgomery Jr, Gale, and Huang 2005; Hashimoto, Elder, and Vitek 2002), independent component analysis (Hyvärinen and Oja 2000; Lemm et al. 2006), linear regression reference (Young et al. 2018), and deep neural networks (Tamada et al. 2020; Zhang and Yu 2018). Of particular note is ERAASR, a technique which uses principal component analysis to exploit the similar structure of the shock artifact sequentially across electrodes, pulses, and then trials (O’Shea and Shenoy 2017). With these approaches, it may be possible to recover neural signal throughout multi-channel stimulation, thereby enabling full band-width recordings in M1 while providing somatosensory feedback via ICMS in S1. Such technology will likely be necessary to accurately decode motor intent as ICMS feedback becomes more complicated.

## Acknowledgements

We would like to thank Tucker Tomlinson and the rest of the Miller Limb Lab for useful discussions that greatly improved this work. This research was funded by National Institute of Neurological Disorders and Stroke Award No. NS095251 and No. F31NS115478, and by National Institute of Health Grant No. T32HD07418. The content is solely the responsibility of the authors and does not necessarily represent the official views of the National Institutes of Health.

## Notes

### Competing Interest Statement

The authors have declared no competing interest.

## References

Allison-Walker, Tim, Maureen A Hagan, Nicholas SC Price, and Yan T Wong. 2021. ‘Microstimulation-evoked neural responses in visual cortex are depth dependent’, Brain Stimulation, 14: 741–50.

Andermann, Mark L, Aaron M Kerlin, and Clay Reid. 2010. ‘Chronic cellular imaging of mouse visual cortex during operant behavior and passive viewing’, Frontiers in cellular neuroscience, 4: 3.

Bensmaia, Sliman J., and Lee E. Miller. 2014. ‘Restoring sensorimotor function through intracortical interfaces: progress and looming challenges’, Nature Reviews Neuroscience, 15: 313–25.

Berg, J. A., Iii J. F. Dammann, F. V. Tenore, G. A. Tabot, J. L. Boback, L. R. Manfredi, M. L. Peterson, K. D. Katyal, M. S. Johannes, A. Makhlin, R. Wilcox, R. K. Franklin, R. J. Vogelstein, N. G. Hatsopoulos, and S. J. Bensmaia. 2013. ‘Behavioral Demonstration of a Somatosensory Neuroprosthesis’, IEEE Transactions on Neural Systems and Rehabilitation Engineering, 21: 500–07.

Brown, Edgar A, James D Ross, Richard A Blum, Yoonkey Nam, Bruce C Wheeler, and Stephen P DeWeerth. 2008. ‘Stimulus-artifact elimination in a multi-electrode system’, IEEE Transactions on Biomedical Circuits and Systems, 2: 10–21.

Butovas, Sergejus, Sheriar G. Hormuzdi, Hannah Monyer, and Cornelius Schwarz. 2006. ‘Effects of Electrically Coupled Inhibitory Networks on Local Neuronal Responses to Intracortical Microstimulation’, Journal of Neurophysiology, 96: 1227–36.

Butovas, Sergejus, and Cornelius Schwarz. 2003. ‘Spatiotemporal Effects of Microstimulation in Rat Neocortex: A Parametric Study Using Multielectrode Recordings’, Journal of Neurophysiology, 90: 3024–39.

Callier, Thierri, Nathan W Brantly, Attilio Caravelli, and Sliman J Bensmaia. 2020. ‘The frequency of cortical microstimulation shapes artificial touch’, Proceedings of the National Academy of Sciences, 117: 1191–200.

Chandrasekaran, Santosh, Stephan Bickel, Jose L Herrero, Joo-won Kim, Noah Markowitz, Elizabeth Espinal, Nikunj A Bhagat, Richard Ramdeo, Junqian Xu, and Matthew F Glasser. 2021. ‘Evoking highly focal percepts in the fingertips through targeted stimulation of sulcal regions of the brain for sensory restoration’, Brain Stimulation, 14: 1184–96.

Chen, Xing, Feng Wang, Eduardo Fernandez, and Pieter R Roelfsema. 2020. ‘Shape perception via a high-channel-count neuroprosthesis in monkey visual cortex’, Science, 370: 1191–96.

Collinger, Jennifer L., Brian Wodlinger, John E. Downey, Wei Wang, Elizabeth C. Tyler-Kabara, Douglas J. Weber, Angus J. C. McMorland, Meel Velliste, Michael L. Boninger, and Andrew B. Schwartz. 2013. ‘High-performance neuroprosthetic control by an individual with tetraplegia’, The Lancet, 381: 557–64.

Dadarlat, Maria C., Joseph E. O’Doherty, and Philip N. Sabes. 2015. ‘A learning-based approach to artificial sensory feedback leads to optimal integration’, Nature Neuroscience, 18: 138–44.

DeMichele, GA, and PR Troyk. 2003. “Stimulus-resistant neural recording amplifier.” In Proceedings of the 25th Annual International Conference of the IEEE Engineering in Medicine and Biology Society (IEEE Cat. No. 03CH37439), 3329–32. IEEE.

Devecioğlu, İsmail, and Burak Güçlü. 2017. ‘Psychophysical correspondence between vibrotactile intensity and intracortical microstimulation for tactile neuroprostheses in rats’, Journal of Neural Engineering, 14: 016010–10.

Dombeck, Daniel A, Anton N Khabbaz, Forrest Collman, Thomas L Adelman, and David W Tank. 2007. ‘Imaging large-scale neural activity with cellular resolution in awake, mobile mice’, Neuron, 56: 43–57.

Epstein, Charles M. 1995. ‘A simple artifact-rejection preamplifier for clinical neurophysiology’, American Journal of EEG Technology, 35: 64–71.

Fifer, Matthew S, David P McMullen, Tessy M Thomas, Luke Osborn, Robert W Nickl, Daniel N Candrea, Eric A Pohlmeyer, Margaret C Thompson, Manuel Anaya, and Wouter Schellekens. 2020. ‘Intracortical microstimulation elicits human fingertip sensations’, medRxiv.

Flesher, S. N., J. L. Collinger, S. T. Foldes, J. M. Weiss, J. E. Downey, E. C. Tyler-Kabara, S. J. Bensmaia, A. B. Schwartz, M. L. Boninger, and R. A. Gaunt. 2016. ‘Intracortical microstimulation of human somatosensory cortex’, Science Translational Medicine, 8: 361ra141–361ra141.

Flesher, Sharlene N, John E Downey, Jeffrey M Weiss, Christopher L Hughes, Angelica J Herrera, Elizabeth C Tyler-Kabara, Michael L Boninger, Jennifer L Collinger, and Robert A Gaunt. 2021. ‘A brain-computer interface that evokes tactile sensations improves robotic arm control’, Science, 372: 831–36.

Freeman, John A. 1971. ‘An electronic stimulus artifact suppressor’, Electroencephalography and clinical neurophysiology, 31: 170–72.

Fridman, Gene Y., Hugh T. Blair, Aaron P. Blaisdell, and Jack W. Judy. 2010. ‘Perceived intensity of somatosensory cortical electrical stimulation’, Experimental Brain Research, 203: 499–515.

Ghez, C., J. Gordon, M. F. Ghilardi, C. N. Christakos, and S. E. Cooper. 1990. ‘Roles of proprioceptive input in the programming of arm trajectories’, Cold Spring Harbor Symposia on Quantitative Biology, 55: 837–47.

Greenberg, David S, Arthur R Houweling, and Jason ND Kerr. 2008. ‘Population imaging of ongoing neuronal activity in the visual cortex of awake rats’, Nature Neuroscience, 11: 749–51.

Gustafsson, B., and E. Jankowska. 1976. ‘Direct and indirect activation of nerve cells by electrical pulses applied extracellularly’, The Journal of Physiology, 258: 33–61.

Hao, Yaoyao, Alexa Riehle, and Thomas G. Brochier. 2016. ‘Mapping Horizontal Spread of Activity in Monkey Motor Cortex Using Single Pulse Microstimulation’, Frontiers in Neural Circuits, 10.

Hashimoto, Takao, Christopher M Elder, and Jerrold L Vitek. 2002. ‘A template subtraction method for stimulus artifact removal in high-frequency deep brain stimulation’, Journal of Neuroscience Methods, 113: 181–86.

Histed, Mark H., Vincent Bonin, and R. Clay Reid. 2009. ‘Direct Activation of Sparse, Distributed Populations of Cortical Neurons by Electrical Microstimulation’, Neuron, 63: 508–22.

Hochberg, Leigh R, Daniel Bacher, Beata Jarosiewicz, Nicolas Y Masse, John D Simeral, Joern Vogel, Sami Haddadin, Jie Liu, Sydney S Cash, and Patrick Van Der Smagt. 2012. ‘Reach and grasp by people with tetraplegia using a neurally controlled robotic arm’, Nature, 485: 372.

Hughes, Christopher L, Sharlene N Flesher, Jeffrey M Weiss, Michael Boninger, Jennifer L Collinger, and Robert A Gaunt. 2021. ‘Perception of microstimulation frequency in human somatosensory cortex’, Elife, 10: e65128.

Hyvärinen, Aapo, and Erkki Oja. 2000. ‘Independent component analysis: algorithms and applications’, Neural networks, 13: 411–30.

Jankowska, E., Y. Padel, and R. Tanaka. 1975. ‘The mode of activation of pyramidal tract cells by intracortical stimuli’, The Journal of Physiology, 249: 617–36.

Jung, Hyunjun, Jintae Kim, and Yoonkey Nam. 2018. ‘Recovery of early neural spikes from stimulation electrodes using a DC-coupled low gain high resolution data acquisition system’, Journal of Neuroscience Methods, 304: 118–25.

Kim, Sungshin, Thierri Callier, Gregg A. Tabot, Robert A. Gaunt, Francesco V. Tenore, and Sliman J. Bensmaia. 2015. ‘Behavioral assessment of sensitivity to intracortical microstimulation of primate somatosensory cortex’, Proceedings of the National Academy of Sciences, 112: 15202–07.

Kumaravelu, Karthik, Joseph Sombeck, Lee E Miller, Sliman J Bensmaia, and Warren M Grill. 2021. ‘Stoney vs. Histed: Quantifying the Spatial Effects of Intracortical Microstimulation’, bioRxiv.

Lemm, Steven, Gabriel Curio, Yevhen Hlushchuk, and K-R Muller. 2006. ‘Enhancing the signal-to-noise ratio of ICA-based extracted ERPs’, IEEE Transactions on Biomedical Engineering, 53: 601–07.

London, B. M., L. R. Jordan, C. R. Jackson, and L. E. Miller. 2008. ‘Electrical Stimulation of the Proprioceptive Cortex (Area 3a) Used to Instruct a Behaving Monkey’, IEEE Transactions on Neural Systems and Rehabilitation Engineering, 16: 32–36.

London, Brian M., and Lee E. Miller. 2012. ‘Responses of somatosensory area 2 neurons to actively and passively generated limb movements’, Journal of Neurophysiology, 109: 1505–13.

Margalit, Shany Nivinsky, and Hamutal Slovin. 2018. ‘Spatio-temporal characteristics of population responses evoked by microstimulation in the barrel cortex’, Scientific Reports, 8.

Maynard, Edwin M, Craig T Nordhausen, and Richard A Normann. 1997. ‘The Utah intracortical electrode array: a recording structure for potential brain-computer interfaces’, Electroencephalography and clinical neurophysiology, 102: 228–39.

McElvain, Lauren E, Martha W Bagnall, Alexandra Sakatos, and Sascha Du Lac. 2010. ‘Bidirectional plasticity gated by hyperpolarization controls the gain of postsynaptic firing responses at central vestibular nerve synapses’, Neuron, 68: 763–75.

Mendrela, Adam E, Jihyun Cho, Jeffrey A Fredenburg, Vivek Nagaraj, Theoden I Netoff, Michael P Flynn, and Euisik Yoon. 2016. ‘A bidirectional neural interface circuit with active stimulation artifact cancellation and cross-channel common-mode noise suppression’, IEEE Journal of Solid-State Circuits, 51: 955–65.

Michelson, Nicholas J., James R. Eles, Alberto L. Vazquez, Kip A. Ludwig, and Takashi D. Y. Kozai. 2018. ‘Calcium activation of cortical neurons by continuous electrical stimulation: Frequency dependence, temporal fidelity, and activation density’, Journal of Neuroscience Research.

Molineux, Michael L, John E McRory, Bruce E McKay, Jawed Hamid, W Hamish Mehaffey, Renata Rehak, Terrance P Snutch, Gerald W Zamponi, and Ray W Turner. 2006. ‘Specific T-type calcium channel isoforms are associated with distinct burst phenotypes in deep cerebellar nuclear neurons’, Proceedings of the National Academy of Sciences, 103: 5555–60.

Montgomery Jr, Erwin B, John T Gale, and He Huang. 2005. ‘Methods for isolating extracellular action potentials and removing stimulus artifacts from microelectrode recordings of neurons requiring minimal operator intervention’, Journal of Neuroscience Methods, 144: 107–25.

Murasugi, Cm, Cd Salzman, and Wt Newsome. 1993. ‘Microstimulation in visual area MT: effects of varying pulse amplitude and frequency’, The Journal of Neuroscience, 13: 1719–29.

Nag, Sudip, Sujit Kumar Sikdar, Nitish Vyomesh Thakor, Valipe Ramgopal Rao, and Dinesh Sharma. 2015. ‘Sensing of stimulus artifact suppressed signals from electrode interfaces’, IEEE Sensors Journal, 15: 3734–42.

Nowak, L. G., and J. Bullier. 1998a. ‘Axons, but not cell bodies, are activated by electrical stimulation in cortical gray matter. I. Evidence from chronaxie measurements’, Experimental Brain Research, 118: 477–88.

Nowak, L. G., and J. Bullier. 1998b. ‘Axons, but not cell bodies, are activated by electrical stimulation in cortical gray matter. II. Evidence from selective inactivation of cell bodies and axon initial segments’, Experimental Brain Research, 118: 489–500.

O’Shea, Daniel J., and Krishna V. Shenoy. 2017. ‘ERAASR: An algorithm for removing electrical stimulation artifacts from multielectrode array recordings’.

O’Doherty, Joseph E., Mikhail A. Lebedev, Peter J. Ifft, Katie Z. Zhuang, Solaiman Shokur, Hannes Bleuler, and Miguel A. L. Nicolelis. 2011. ‘Active tactile exploration using a brain–machine–brain interface’, Nature, 479: 228–31.

Otto, Kevin J., Patrick J. Rousche, and Daryl R. Kipke. 2005. ‘Cortical microstimulation in auditory cortex of rat elicits best-frequency dependent behaviors’, Journal of Neural Engineering, 2: 42–51.

Overstreet, C. K., J. D. Klein, and S. I. Helms Tillery. 2013. ‘Computational modeling of direct neuronal recruitment during intracortical microstimulation in somatosensory cortex’, Journal of Neural Engineering, 10: 066016.

Rolston, JD, RE Gross, and SM Potter. 2009. ‘A low-cost multielectrode system for data acquisition and real-time processing with rapid recovery from stimulation artifacts’, Frontiers in Neuroengineering, 2: 1–17.

Romo, Ranulfo, Adrián Hernández, Anótonio Zainos, and Emilio Salinas. 1998. ‘Somatosensory discrimination based on cortical microstimulation’, Nature, 392: 387–90.

Romo, Ranulfo, Adrián Hernández, Antonio Zainos, Carlos D. Brody, and Luis Lemus. 2000. ‘Sensing without Touching: Psychophysical Performance Based on Cortical Microstimulation’, Neuron, 26: 273–78.

Sainburg, R. L., M. F. Ghilardi, H. Poizner, and C. Ghez. 1995. ‘Control of limb dynamics in normal subjects and patients without proprioception’, Journal of Neurophysiology, 73: 820–35.

Salas, Michelle Armenta, Luke Bashford, Spencer Kellis, Matiar Jafari, HyeongChan Jo, Daniel Kramer, Kathleen Shanfield, Kelsie Pejsa, Brian Lee, Charles Y. Liu, and Richard A. Andersen. 2018. ‘Proprioceptive and cutaneous sensations in humans elicited by intracortical microstimulation’, Elife, 7.

Salzman, C Daniel, Chieko M Murasugi, Kenneth H Britten, and William T Newsome. 1992. ‘Microstimulation in visual area MT: effects on direction discrimination performance’, Journal of Neuroscience, 12: 2331–55.

Sombeck, Joseph T, and Lee E Miller. 2019. ‘Short reaction times in response to multi-electrode intracortical microstimulation may provide a basis for rapid movement-related feedback’, Journal of Neural Engineering, 17: 016013.

Stieger, Kevin C, James R Eles, Kip A Ludwig, and Takashi DY Kozai. 2020. ‘In vivo microstimulation with cathodic and anodic asymmetric waveforms modulates spatiotemporal calcium dynamics in cortical neuropil and pyramidal neurons of male mice’, Journal of Neuroscience Research, 98: 2072–95.

Stoney, S. D., W. D. Thompson, and H. Asanuma. 1968. ‘Excitation of pyramidal tract cells by intracortical microstimulation: effective extent of stimulating current’, Journal of Neurophysiology, 31: 659–69.

Swadlow, Harvey A. 1990. ‘Efferent neurons and suspected interneurons in S-1 forelimb representation of the awake rabbit: receptive fields and axonal properties’, Journal of Neurophysiology, 63: 1477–98.

Tabot, Gregg A., John F. Dammann, Joshua A. Berg, Francesco V. Tenore, Jessica L. Boback, R. Jacob Vogelstein, and Sliman J. Bensmaia. 2013. ‘Restoring the sense of touch with a prosthetic hand through a brain interface’, Proceedings of the National Academy of Sciences of the United States of America, 110: 18279–84.

Tamada, Daiki, Marie-Luise Kromrey, Shintaro Ichikawa, Hiroshi Onishi, and Utaroh Motosugi. 2020. ‘Motion artifact reduction using a convolutional neural network for dynamic contrast enhanced MR imaging of the liver’, Magnetic Resonance in Medical Sciences, 19: 64.

Tanaka, Yuta, Tomohiro Nomoto, Takuya Shiki, Yuya Sakata, Yoshinori Shimada, Yuki Hayashida, and Tetsuya Yagi. 2019. ‘Focal activation of neuronal circuits induced by microstimulation in the visual cortex’, Journal of Neural Engineering, 16: 036007.

Tehovnik, E. J., A. S. Tolias, F. Sultan, W. M. Slocum, and N. K. Logothetis. 2006. ‘Direct and Indirect Activation of Cortical Neurons by Electrical Microstimulation’, Journal of Neurophysiology, 96: 512–21.

Tehovnik, Edward J., and Warren M. Slocum. 2013. ‘Two-photon imaging and the activation of cortical neurons’, Neuroscience, 245: 12–25.

Tomlinson, Tucker, and Lee E. Miller. 2016. ‘Toward a Proprioceptive Neural Interface that Mimics Natural Cortical Activity.’ in Jozsef Laczko and Mark L. Latash (eds.), Progress in Motor Control (Springer International Publishing: Cham).

Venkatraman, Subramaniam, Ken Elkabany, John D Long, Yimin Yao, and Jose M Carmena. 2008. ‘A system for neural recording and closed-loop intracortical microstimulation in awake rodents’, IEEE Transactions on Biomedical Engineering, 56: 15–22.

Weiss, Jeffrey M, Sharlene N Flesher, Robert Franklin, Jennifer L Collinger, and Robert A Gaunt. 2018. ‘Artifact-free recordings in human bidirectional brain–computer interfaces’, Journal of Neural Engineering, 16: 016002.

Wodlinger, B, JE Downey, EC Tyler-Kabara, AB Schwartz, ML Boninger, and JL Collinger. 2014. ‘Ten-dimensional anthropomorphic arm control in a human brain− machine interface: difficulties, solutions, and limitations’, Journal of Neural Engineering, 12: 016011.

Young, D, F Willett, WD Memberg, B Murphy, B Walter, J Sweet, J Miller, LR Hochberg, RF Kirsch, and AB Ajiboye. 2018. ‘Signal processing methods for reducing artifacts in microelectrode brain recordings caused by functional electrical stimulation’, Journal of Neural Engineering, 15: 026014.

Zhang, Yanbo, and Hengyong Yu. 2018. ‘Convolutional neural network based metal artifact reduction in x-ray computed tomography’, IEEE transactions on medical imaging, 37: 1370–81.

Zhou, Andy, Benjamin C. Johnson, and Rikky Muller. 2018. ‘Toward true closed-loop neuromodulation: artifact-free recording during stimulation’, Current Opinion in Neurobiology, 50: 119–27.

