## Supplementary Information for "Characterizing the short-latency evoked response to intracortical microstimulation across a multi-electrode array"

### Supplementary Materials

#### Technical description of the rapid-recovery amplifier

We aimed to develop an amplifier which can record neural activity ~1 ms after stimulation offset. To do so, the stimulus pulses should not overdrive the amplifier nor saturate the filters, particularly high-pass filters, which by design have a long recovery time constant. The amplifier schematics are shown in Figure S1. The amplifier implements a single-ended configuration with three stages; the first stage uses the OPA140DGK (Texas Instruments, Dallas, TX) operational amplifier (op-amp) and the second and third stages use OPA2277U-EP (Texas Instruments). The first (input) stage and the second stage include high-pass filters and gain compression; the third stage is a linear gain stage.

All stages are supplied with ±15 V to prevent output saturation and input current shunting. If the op-amp input were to exceed the power supply rails, the input signal would be clamped to ±15 V due to the electrostatic discharge protection of the chip, resulting in an undesirable transient drop in the input impedance. As a consequence, the input stage might shunt a part of the stimulation current, reducing the current left for stimulation if the stimulator and amplifier share the same electrode. If stimulation uses a separate electrode, the drop in impedance might inject a significant current through the recording electrode.

We used an ac-coupling input capacitance of 1 nF to balance several design objectives. A large capacitance value is advantageous as it reduces the impedance driving the input stage, extends the low-frequency corner, and reduces the residual voltage on the capacitor after a large input artifact, which could saturate the input stage. On the other hand, a small capacitance value is required to limit the charge injected into the brain in case of an op-amp failure, which may short the op-amp input to its ± 15 V supply. The ac-coupling capacitance is biased to ground with a 500 MΩ resistor, which dominates the input impedance. As the input impedance forms a voltage divider together with the electrode and the source's equivalent impedance, the input impedance should be significantly larger than the total impedance driving the amplifier input. The high ac-coupling capacitance and high impedance leads to a long time constant of the first stage (500 ms), which is eliminated by faster (1 ms) ac-coupling dynamics in the second stage. To allow a high input impedance without a large dc voltage offset caused by the op-amp input bias current, the feedback network of the first stage is ac-coupled to ground through a 570 nF capacitor, reducing the dc gain to unity. This capacitance is larger than the ac-coupling input capacitance, but does not affect the maximum injected charge in case of a failure of an integrated circuit.

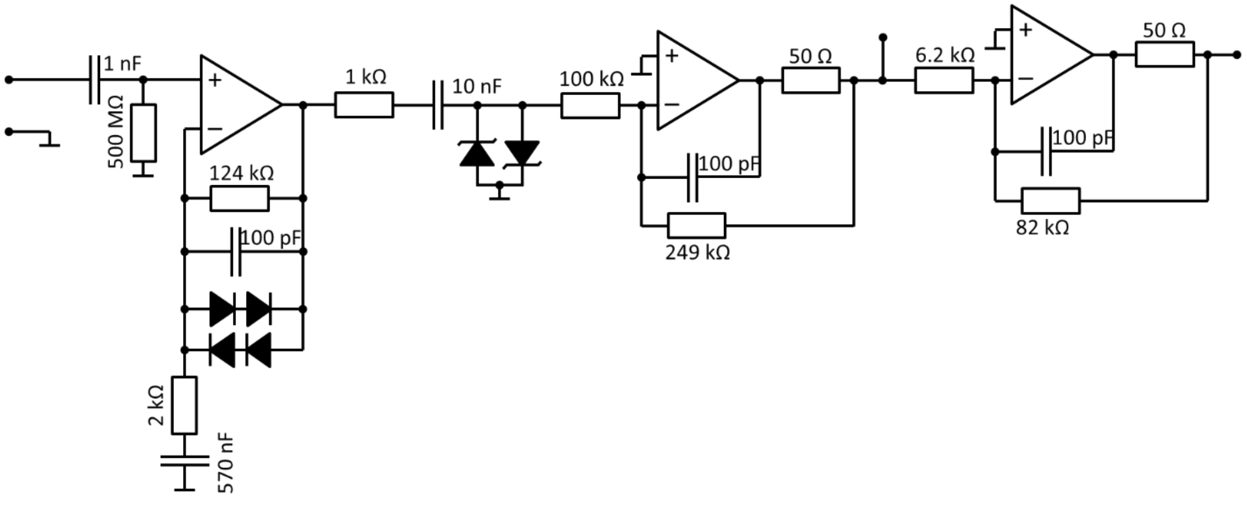


**Fig. S1.** Circuit diagram of the rapid-recovery amplifier

Saturation of subsequent stages during a stimulus pulse is avoided by nonlinear gain, which compresses large dynamic swings. We implemented a feedback loop in the first stage which reduces the gain to unity when the output exceeds the input by approximately 1.3 V, corresponding the total forward voltage drop of the two series diodes in parallel with the feedback resistor and capacitor. The input to the second amplification stage is ac-coupled with a fast time constant (1 ms) and is clamped by a pair of Schottky diodes limiting the gain for large signals (Mueller et al. 2014). The rest of the second stage and the third stage are linear and provide additional gain for the neural signals.

#### Finding peaks in the evoked response

Spikes were often evoked at multiple, consistent latencies following single pulses of stimulation. Fig. S2 shows the response of an example neuron to repeated stimulation at 50 µA (top, same format as Fig. 3). We grouped spikes based on their response latency in each condition. To do so, we first convolved the spike train with a non-causal Gaussian kernel (width = 0.2 ms) (Fig. S2 bottom, black trace). We found peaks in this average with MATLAB's *findpeaks* algorithm. This algorithm computes the "prominence" of every local maximum by first drawing horizontal lines before and after each maximum until the lines either cross the signal or reach the end of the signal (horizontal lines in Fig. S2). Then, the minimum of the signal is found within both intervals, one before and one after the local maximum. Prominence is the difference between the local maximum and the highest of these two minimums (vertical lines in Fig. S2). If the prominence for a local maximum is sufficiently large (above 1.0 in our case), then the maximum is labelled as a peak. Peaks that were found with this algorithm are marked with green circles and a subset of local maxima that were rejected are marked with red squares (Fig. S2). This algorithm also measures the width of the peak, defined as the time between points where the descending signal intercepts half of the peak prominence, shown as blue lines within each peak.

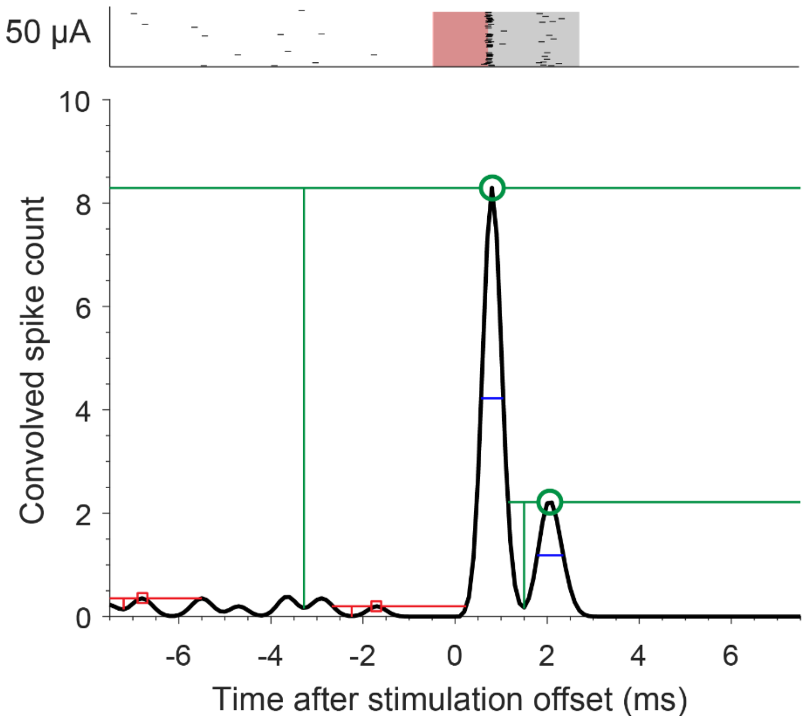


**Fig. S2.** Example prominence calculation and resulting peaks using *findpeaks* algorithm.

#### Evoked response on the stimulated channel during intermittent stimulation

Since the response on the stimulated channel decays rapidly, we wondered whether using intermittent stimulation to allow neurons time to recover would reduce the decay rate. As opposed to stimulation at a fixed frequency, we turned stimulation on and off throughout the train. We varied both the duty cycle (33%, 50%, and 67%) and the duration (50, 100, and 200 ms) of the stimulation. A duty cycle of 33% and duration of 100 ms means that stimulation was on for 100 ms, then off for 200 ms throughout a 4 s long train. We delivered 40 µA pulses at either 131 or 179 Hz. When stimulating continuously, we used a frequency roughly equal to the duty cycle times the stimulation frequency to match the number of pulses. We could not use the exact equivalent frequencies because we were limited in the frequencies we could stimulate with while simultaneously recording on the stimulated electrode.

Fig. S3a shows the average activity evoked in an example neuron recorded on the stimulated electrode for six of the nine intermittent stimulation conditions at 179 Hz. The decay rates for neurons recorded on the stimulated electrode are summarized for both frequencies (Fig. S3b,c). We fit a linear model to predict decay rate from duty cycle and duration ($F(28,241) = 4.78$, $p<<0.001$). Increasing the duty cycle significantly increased the decay rate regardless of stimulation frequency (duty cycle factor $F(1,241) = 24.07$, $p = 1.7\times10^{-6}$), as increasing this parameter also increased the amount of charge delivered during the 4-s train. Increasing duration also significantly increased the decay rate ($F(1,241) = 5.6$, $p = 0.019$), although this effect was an order of magnitude smaller than the duty cycle parameter, possibly because increasing duration does not increase the amount of charge. Aggregated across durations and neurons, intermittent stimulation did not significantly change the decay rate for matched numbers of pulses and total charge (Wilcoxon rank-sum test, $p > 0.05$ for all), indicating that the effect intermittent stimulation has on decay rate is similar to simply reducing the mean stimulus frequency or number of stimulus pulses.

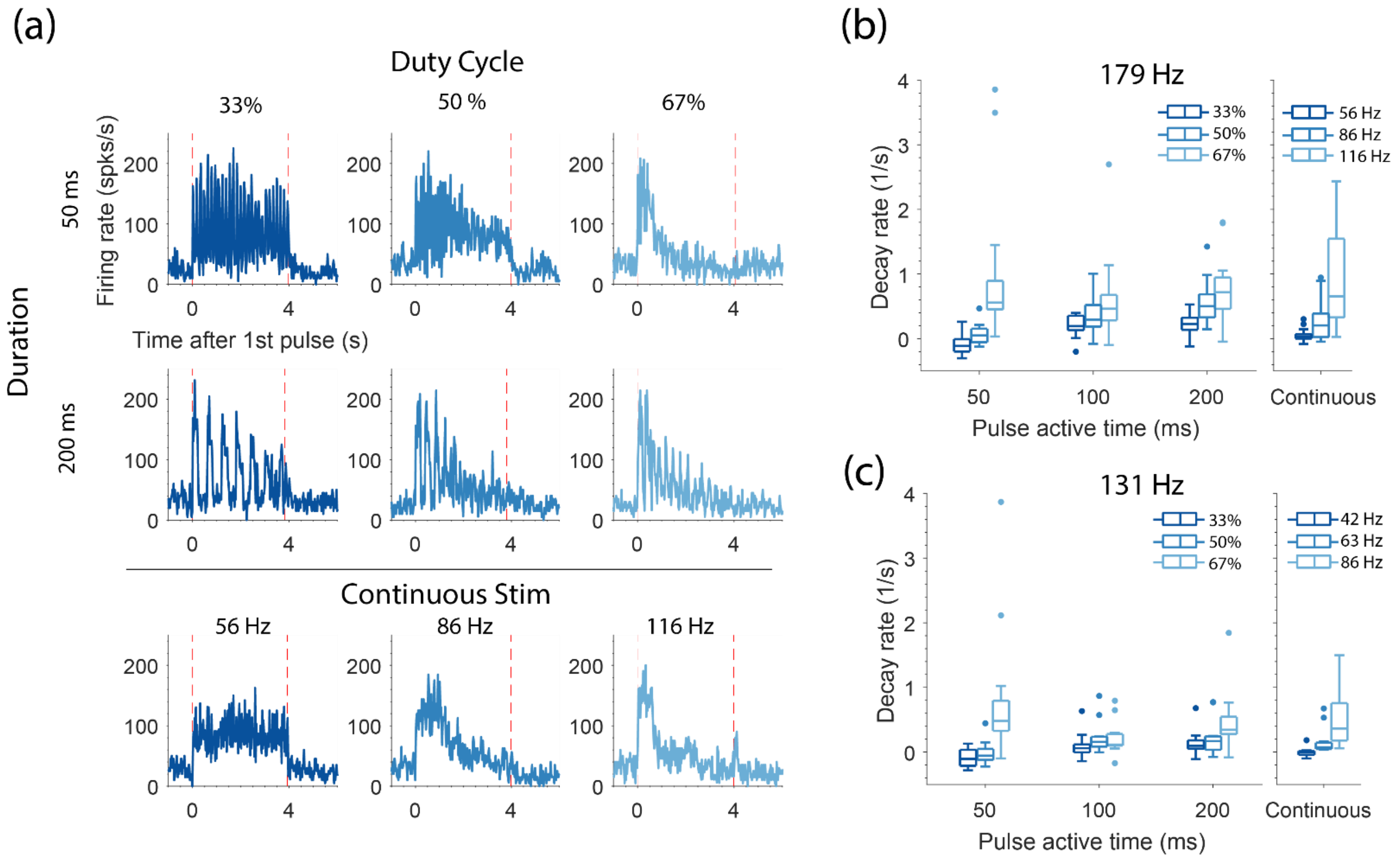


**Fig. S3.** Evoked response on the stimulated channel during intermittent stimulation. (a) The mean firing rate across stimulation trials for an example neuron recorded on the stimulated electrode for different duty cycles (columns) and durations (rows). Stimulation frequency was 179 Hz. The mean firing rate is also shown during continuous stimulation at control frequencies. (b) The decay rates across neurons and conditions when stimulating at 179 Hz. The decay rates across the same neurons are shown during continuous stimulation. Points indicate outliers. (c) Same as (b) except when stimulating at 131 Hz.
